# Network-informed discovery of multidrug combinations for ERα+/HER2-/PI3Kα-mutant breast cancer

**DOI:** 10.1101/2022.08.30.505871

**Authors:** Dina Hany, Marloes Zoetemelk, Kaushik Bhattacharya, Patrycja Nowak-Sliwinska, Didier Picard

## Abstract

Breast cancer is a persistent threat to women worldwide. A large proportion of breast cancers are dependent on estrogen receptor α (ERα) for tumor progression. Therefore, targeting ERα with antagonists, such as tamoxifen, remains standard therapy for ERα+ breast cancer. The clinical benefits of monotherapy are often counterbalanced by off-target toxicity and development of resistance. Combinations of more than two drugs might be of great therapeutic value to prevent resistance, and to reduce doses, and hence, toxicity. We mined data from the literature and public repositories to construct a network of potential drug targets for synergistic multidrug combinations. With 9 drugs, we performed a phenotypic combinatorial screen with ERα+ breast cancer cell lines. We identified two optimized low-dose combinations of 3 and 4 drugs of high therapeutic relevance to the frequent ERα+/HER2-/PI3Kα-mutant subtype of breast cancer. Moreover, we validated the efficacy of the combinations in tamoxifen-resistant cell lines, patient-derived organoids, and xenograft experiments. Thus, we propose multidrug combinations that have the potential to overcome the standard issues of current monotherapies.

## Introduction

Breast cancer is the most prevalent type of cancer among women worldwide (Sung *et al*, 2021). Therapeutic strategies for the treatment of breast cancer are guided by clinical features and the histological and molecular subtyping of the disease. Estrogen receptor α (ERα) is expressed in more than two thirds of breast tumors and marks a favorable prognosis of the disease (Shiino *et al*, 2016). In these tumors, ERα represents a focal point of cellular signaling that regulates cell growth, proliferation, survival, invasion, and metastasis. ERα is a multidomain transcription factor of the nuclear receptor superfamily. The binding of estrogens, such as 17β-estradiol (E2), to ERα initiates its dimerization and activation of extensive genomic and non-genomic signaling. Recruiting several coregulators, activated ERα binds either directly to specific DNA sequences known as estrogen response elements (EREs) or indirectly to the response elements of other transcription factors, including nuclear factor kappa B (NFκB), which collectively leads to the transcriptional regulation of a wide spectrum of genes (Klinge, 2001; Marino *et al*, 2006; Hall *et al*, 2002). Apart from ERα-mediated genomic effects, activation of a subset of membrane-associated ERα molecules can trigger the rapid activation of cell signaling cascades, including those involving phosphatidylinositol 3-kinase (PI3K), protein kinase B (hereafter referred to as Akt), mammalian target of rapamycin (mTOR), and mitogen-activated protein kinases (MAPKs), which eventually leads to cell proliferation, differentiation, and survival (Levin, 2005). In addition, ERα can be activated in an estrogen-independent manner by a large panel of factors, including epidermal growth factor (EGF) (Ignar-Trowbridge *et al*, 1992; Ignar-Trowbridge *et al*, 1993), insulin and insulin-like growth factors (IGF) (Newton *et al*, 1994), PI3K (Campbell *et al*, 2001), Akt (Campbell *et al*, 2001; Martin *et al*, 2000), and hypoxia-inducible factor 1 α (HIF1α) (Cho *et al*, 2006; Bennesch & Picard, 2015) (see detailed list at https://www.picard.ch/downloads/Factors.pdf). Through various domains, ERα can interact with a plethora of proteins that regulate its activity, such as the MAPKs ERK1/ERK2 (ERK1/2) (Chen *et al*, 2002; Kato *et al*, 1995; Kim *et al*, 2021; Vafeiadou *et al*, 2022; Li *et al*, 2012; Mueller *et al*, 2000; Bunone *et al*, 1996), mTOR (Alayev *et al*, 2016), cyclin-dependent kinase inhibitor 1 (p21) (Fritah *et al*, 2005; Redeuilh *et al*, 2002; Abukhdeir *et al*, 2008), and poly (ADP-ribose) polymerase 1 (PARP1) (Schiewer & Knudsen, 2014; Zhang *et al*, 2013; Pulliam *et al*, 2019; Gadad *et al*, 2021) (see the updated list of ERα interactors at https://www.picard.ch/downloads).

Targeting ERα with endocrine therapy has been the gold standard for decades (Aggelis & Johnston, 2019). This includes the use of selective ERα modulators, such as tamoxifen and raloxifene, selective ERα degraders, such as fulvestrant, and aromatase inhibitors to deprive ERα of estrogens. Tamoxifen is the first-line drug for endocrine therapy used in the treatment (Jordan, 2003) and prevention (Cuzick *et al*, 2003) of ERα+ breast cancer; it markedly increases patient survival and is relatively well tolerated (Jordan, 2003; Jankowitz & Davidson, 2013). However, around 40% of patients with localized and metastatic breast cancer develop endocrine resistance (Hultsch *et al*, 2018). Resistance to tamoxifen can be attributed to multiple mechanisms, including loss of ERα expression or function due to genetic mutations, alterations in the levels of co-regulatory factors (Osborne *et al*, 2003), upregulation of oncogenic signal transduction pathways, and ligand-independent activation of ERα (Kato *et al*, 1995; Shim *et al*, 2000; Coutts & Murphy, 1998; Gee *et al*, 2001; Clark *et al*, 2002; Gutierrez *et al*, 2005; Bennesch & Picard, 2015). For example, ERK1/2 play a critical role in breast cancer and their hyperactivation is associated with tamoxifen resistance and poor prognosis (Gee *et al*, 2001; Coutts & Murphy, 1998; Shim *et al*, 2000; Chen *et al*, 2002; Kato *et al*, 1995; Kim *et al*, 2021; Vafeiadou *et al*, 2022; Li *et al*, 2012; Mueller *et al*, 2000; Bunone *et al*, 1996). ERK1/2 can stimulate cell growth independently of ERα or through signaling crosstalk with ERα. ERK1/2 phosphorylates ERα at S118, which promotes ligand-independent activity (Bunone *et al*, 1996; Kato *et al*, 1995). Another example is the PI3K/Akt/mTOR signaling pathway. Activating mutations in the p110α subunit of PI3K (PI3Kα) encoded by the gene *PIK3CA* are the most common mutations in ERα+/HER2-breast cancer; they occur in approximately 40% of patients and are associated with poor prognosis and resistance to both chemotherapy and endocrine therapy (Fusco *et al*, 2021; Wu *et al*, 2005; Mosele *et al*, 2020; Bosch *et al*, 2015; Martínez-Saéz *et al*, 2020). Activation of PI3Kα leads to the activation of Akt and mTOR, both of which can phosphorylate ERα at sites, which stimulate its transcriptional activity, eventually leading to tamoxifen resistance (Campbell *et al*, 2001; Bennesch & Picard, 2015; de Leeuw *et al*, 2011; Alayev *et al*, 2016). Moreover, downstream of mTOR is NFκB, whose activation is associated with estrogen-independent growth and endocrine therapy resistance (Zhou *et al*, 2005; deGraffenried *et al*, 2004; Nakshatri *et al*, 1997).

The rational use of cancer monotherapy is limited by the activation of bypass mechanisms, feedback or feedforward regulatory loops, and signaling crosstalk that eventually lead to drug resistance (Iadevaia *et al*, 2010). Drug combinations involve the multi-dimensional targeting of biological networks (Keith *et al*, 2005; Weiss *et al*, 2019), which increases the efficacy and reduces the likelihood of developing resistance to the simultaneously used drugs (Bozic *et al*, 2013). For ERα+ breast cancer, combinations of 2 drugs commonly available in the clinic can be highly efficacious and synergistic, and yet, they can be associated with toxicity due to the relatively high doses used to attain the desired efficacy. Based on the results of several randomized phase III clinical trials (Lorfida *et al*, 2020), combinations of the ERα degrader fulvestrant with CDK4/6 inhibitors, such as palbociclib, ribociclib, or abemaciclib, have recently been approved for the management of ERα+/HER2-metastatic breast cancer. Moreover, a randomized clinical trial that combined fulvestrant with the PI3Kα inhibitor alpelisib led to the FDA approval of this 2-drug combination for postmenopausal women with ERα+/HER2-/ PI3Kα-mutant advanced or metastatic breast cancer (Narayan *et al*, 2021). However, here too, patients eventually acquire drug resistance (Turner *et al*, 2019; O’Leary *et al*, 2018b; O’Leary *et al*, 2018a; Razavi *et al*, 2020). In addition, these combinations require relatively high doses, which are associated with side effects (Rugo *et al*, 2020; Lorfida *et al*, 2020). Therefore, higher-order combinations of more than 2 drugs at very low doses are of increasing clinical interest (Rausch *et al*, 2020a; Nowak-Sliwinska *et al*, 2016; Pascual *et al*, 2021; Weiss *et al*, 2019; Weiss *et al*, 2015a).

Optimization of drug combinations involves a multistep process. It includes the selection of candidate drug targets and the relevant inhibitors or activators, selection of the optimal dose ratios, design of combinatorial drug matrices, identification of synergistic and antagonistic drug interactions, and evaluation of potential toxicity (Weiss *et al*, 2019). Prior knowledge, clinical experience, or guesswork usually influence the initial selection of drug targets (Decker & Sausville, 2005; Cheng *et al*, 2019), even though mechanism-driven and network-based approaches can be of greater predictive value (Decker & Sausville, 2005; Lehár *et al*, 2007; Iadevaia *et al*, 2010; Sun *et al*, 2015; Cheng *et al*, 2019). However, this is not trivial either because of the size, dynamics, and complexity of biological networks in cancer (Sun *et al*, 2015).

Experimental testing of drug combinations can be done by “exhaustive” or “efficient” approaches (Lehár *et al*, 2007). Exhaustive approaches usually involve testing all possible combinations of the drugs being tested (Tan *et al*, 2012; Lehár *et al*, 2007). In contrast, an efficient approach, such as the “streamlined feedback system control” (s-FSC) method, which is a rapid iterative approach, can reduce the number of tested drug combinations to 0.1% of the total possibilities (Ding *et al*, 2012; Tsutsui *et al*, 2011; Honda *et al*, 2013; Weiss *et al*, 2015b; Wang *et al*, 2015; Ding *et al*, 2014; Nowak-Sliwinska *et al*, 2016). Through only few rounds of experimental assays, regression modeling identifies drugs that lack efficacy or show antagonistic interactions in combinations, and it suggests changes in dose ratios to enhance the interaction between the drugs (Weiss *et al*, 2015a).

The “Therapeutically Guided Multidrug Optimization” (TGMO) method was subsequently developed based on the s-FSC method. It has the advantage of testing drug combinations in malignant and non-malignant cell lines simultaneously and uses therapeutically relevant drug doses, which are below clinically reported ones. This enables the identification of low-dose synergistic drug combinations with favorable safety profiles profiles (Weiss *et al*, 2019; Zoetemelk *et al*, 2020; Rausch *et al*, 2020a; Rausch *et al*, 2020b; Rausch *et al*, 2021).

Owing to the limitations of the currently approved therapies for ERα+ breast cancer, including the development of resistance and drug-related toxicity, there is an urgent need for new and improved drug cocktails. In the current study, we constructed a network of potential drug targets for combination therapy in ERα+ breast cancer by *in silico* data integration. Moreover, using the *in vitro* TGMO-based screening platform, we identified optimized low-dose synergistic drug combinations of 3 and 4 drugs with high efficacy in the tamoxifen-sensitive and -resistant subtypes of ERα+/HER2-/PI3Kα-mutant breast cancer, and with reduced toxicity in a non-cancerous breast epithelial cell line.

## Results

### *In silico* database screen identifies potential drug targets in ERα+ breast cancer and endocrine resistance

The identification of drug targets, which could be targeted synergistically with ERα, can form the basis for the discovery of efficient drug combinations. We employed an *in silico* approach to integrate data from multiple sources, including the scientific literature and public databases. Using the software Cytoscape, we constructed a protein-protein interaction (PPI) network composed of ERα as a central node with its 322 protein interactors and secondary interactions among them (Han *et al*, 2004; Shannon *et al*, 2003) (Fig. 1A and Data S1). The literature-curated list of experimentally validated ERα interactors (Data S2) was from the continuously updated list at http://www.picard.ch/downloads (previously mentioned in Uversky *et al*, 2005; Dunker *et al*, 2015). Since a highly connected protein in a PPI network, known as a “hub protein”, is more likely to be of biological importance than a less connected one (Han *et al*, 2004), we analyzed multiple topological centrality measures, including average shortest path length, number of directed edges, stress, and radiality using the “NetworkAnalyzer” tool available for Cytoscape (Assenov *et al*, 2008; Doncheva *et al*, 2012) (Data S3). From this analysis, we extracted the top 50 protein interactors in the network, including ERα (ESR1), TP53, JUN, EP300, and CREBBP (CBP) (Fig. 1A, B). Next, we explored the gene expression profiles of ERα+ breast cancer clinical biopsies for co-expression with ERα. This yielded 224 and 213 genes from Oncomine and GOBO, respectively (Data S4). The top 50 genes included *GATA3*, *CA12*, *TBC1D9*, and *FOXA1* (Fig. 1C). Then, we mined the mutational landscape of ERα+ breast cancer, and extracted the most frequently mutated genes, including *PIK3CA*, *TP53*, *ESR1*, *CDH1*, and *MAP3K1* (Fig. 1D, E, and Fig. S1A, B).

**Fig. 1.**
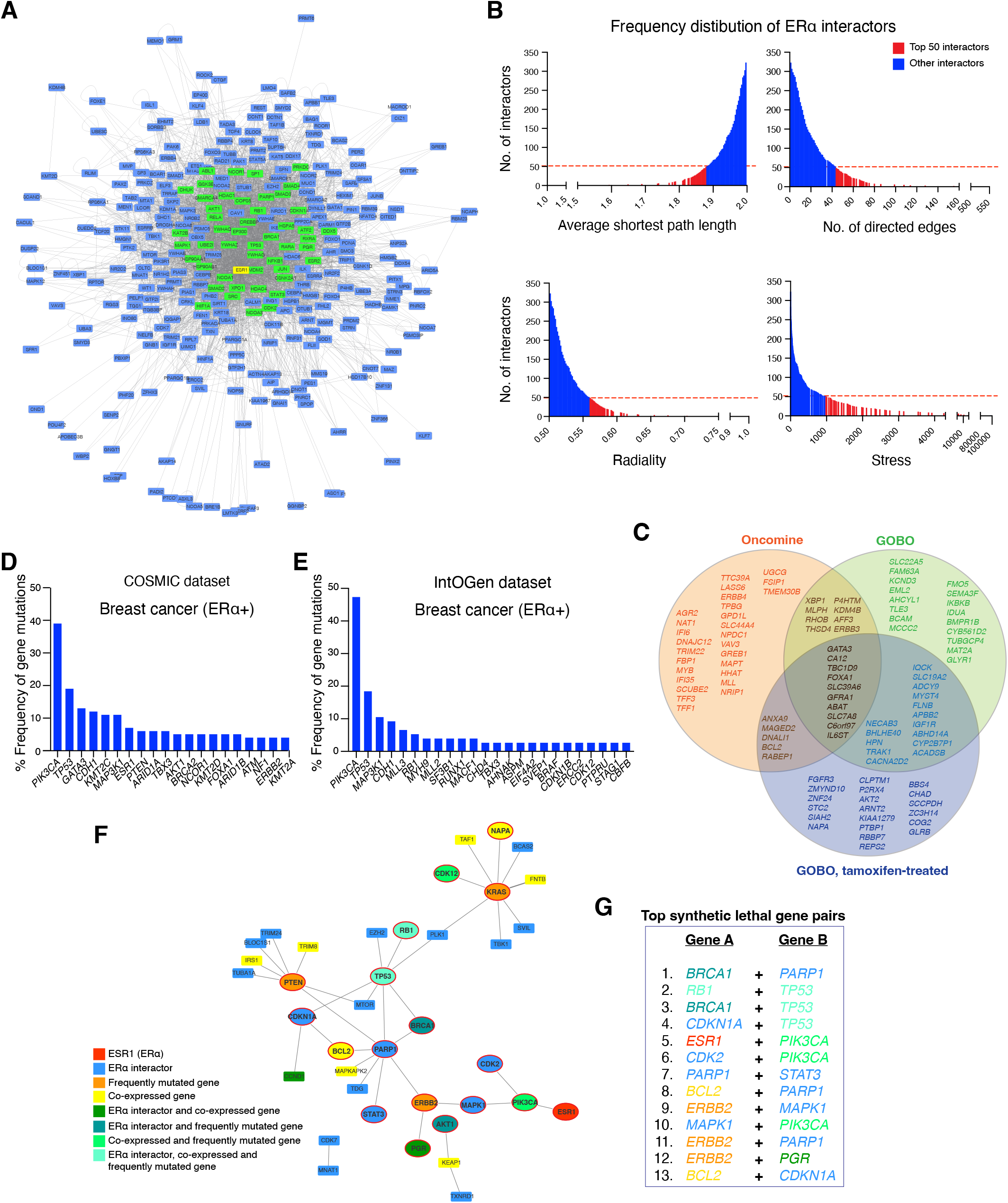
*In silico* database screen identifies potential drug targets in ERα+ breast cancer and endocrine resistance. **(A)** ERα-related interactome composed of 322 ERα interactors (represented as nodes) and both primary and secondary interactions within the PPI network shown as connecting lines (edges) in gray. The top 50 interactors with the highest connectivity and centrality measures are highlighted in green. The PPI network image is zoomable and the original Cytoscape file is in Data S1. **(B)** Frequency distribution histograms of ERα interactors according to node centrality measures, that is average shortest path length (top left), number of directed edges (top right), radiality (bottom left), and stress (bottom right). Bars that correspond to the top 50 ERα interactors are highlighted in red. **(C)** Venn diagram that represents 3 datasets of genes co-expressed with ERα in ERα+ breast cancer. The datasets include the top 50 genes from Oncomine, GOBO, and the GOBO dataset of tamoxifen-treated breast tumors. **(D,E)** Frequency of gene mutations in ERα+ breast cancer (in %) obtained from the databases COSMIC (panel D) and IntOGen (panel E). **(F)** An integrated network of SL between ERα interactors, genes co-expressed with ERα, and frequently mutated genes in ERα+ breast cancer. Nodes represent genes/proteins and edges represent SL relationships between the connected genes. **(G)** A list of the top SL gene pairs ranked based on SL score.

Finally, we included the concept of synthetic lethality (SL) (Hartwell *et al*, 1997). SL indicates that a loss of a gene pair results in lethality while the single gene loss in a particular gene pair does not, SL might therefore suggest effective drug combinations (Heinzel *et al*, 2019). We extracted information about SL between genes in breast cancer from the Synthetic Lethality Database (SynLethDB) (Guo *et al*, 2016) and the siRNA-validated list of cancer-driving synthetically lethal gene pairs (Ye *et al*, 2016). In both datasets, we gave priority to experimentally validated synthetically lethal gene pairs rather than predicted ones, and we established a ranked list of synthetically lethal gene pairs in breast cancer that could be visualized as a network of SL using Cytoscape (Fig. S1C and Data S5). Then, we integrated all extracted datasets (ERα interactome, gene expression profiles, mutational landscape, and synthetic lethal gene pairs) into one network of 38 synthetically lethal gene pairs that contains 23 ERα interactors, 14 genes co-expressed with ERα, and 7 frequently mutated genes in ERα+ breast cancer (Fig. 1F). From this integrated network, we selected the top synthetically lethal gene pairs to search for relevant small molecule inhibitors or activators (Fig. 1G). We prioritized FDA-approved drugs and those in clinical trials. We also considered drug-target selectivity/specificity and drug-drug interactions as important criteria. In the end, our list included nine drugs (Table S1). Signaling pathways involving the selected drug targets and their potential crosstalk are illustrated in Fig. S1D.

### *In vitro* TGMO-based screen identifies synergistic 2-, 3-, and 4-drug combinations for ERα+ breast cancer cell lines

Our goal was to use cell lines to screen for synergistic drug combinations at low doses with maximized efficacy on breast cancer and minimized toxicity on non-cancerous cells. Considering the substantial number of combinations that would be possible with 9 drugs, each added at 2 different concentrations in addition to the vehicle only condition, we resorted to the TGMO-based screening platform (Nowak-Sliwinska *et al*, 2016; Rausch *et al*, 2020a; Weiss *et al*, 2019). It enabled us to identify synergistic and relatively safe drug combinations with limited numbers of experimental tests. Using the suggested design, we tested 91 combinations instead of 3^9^ combinations (i.e. 19,683 assays). The approach is based on a phenotypic assay of the cellular ATP levels, which reflect the metabolic activity, and hence, the viability of cells. We initially screened drug combinations using the three ERα+ breast cancer cell lines, MCF7 (tamoxifen-sensitive), MCF7-V (a more tamoxifen-tolerant variant of MCF7) (Vafeiadou *et al*, 2022; Mohammadi Ghahhari *et al*, 2022), and MCF7/LCC2 (tamoxifen-resistant) (Brünner *et al*, 1993). In parallel, we used MCF10A, a non-malignant breast epithelial cell line, to monitor the relative toxicity, and to calculate the therapeutic window (TW), which represents the difference in efficacy between cancerous and non-cancerous cells.

We generated dose-response curves for each drug with the afore-mentioned cell lines, and with the two additional non-malignant cell lines RPE1 (a retinal pigment epithelial cell line immortalized with telomerase), and CCD-18Co (normal colon cell line). 4-hydroxytamoxifen (OHT, an active metabolite of tamoxifen), alpelisib, the p21 (CDKN1A) inhibitor UC2288 (Wettersten *et al*, 2013), and the STAT3 inhibitor C188-9 (Chen *et al*, 2021a) showed a wide TW, and hence, more selectivity for cancer cells and less toxicity as monotherapies (Fig. S2). Based on the dose-response curves, we selected two doses for each drug, where the higher of the two doses corresponds to the concentration required for a 20% inhibition (IC20), and the lower dose is usually half of that (Table S1). Of note, the selected low doses are below the corresponding maximum tolerated doses and the maximum drug plasma concentrations that were reported in clinical or preclinical studies. For each drug, we tested 3 conditions, including the two selected doses and a vehicle control without the drug. We allocated the 3 conditions in the combinatorial design that contains the 9 drugs (Fig. S3). Next, we performed second-order or third-order linear regression analyses of the calculated % ATP levels of the tested drug combinations using the software MATLAB (Weiss *et al*, 2015a). The estimated regression coefficients obtained from the regression models quantify the contribution of each drug alone (single-drug linear effect and single-drug quadratic effect) and the overall contributions of 2-drug combinations (2-drug interaction), or 3-drug combinations (3-drug interaction in case of a third-order linear regression analysis). Similarly, we estimated the regression coefficient of the TW (Fig. 2A). Negative coefficients indicate a synergistic contribution, while positive coefficients indicate an antagonistic contribution of the drug or its combinations (Fig. 2A). Therefore, an ideal drug combination would have a negative regression coefficient in cancer cells and a positive regression coefficient for the TW (Fig. 2A).

**Fig. 2.**
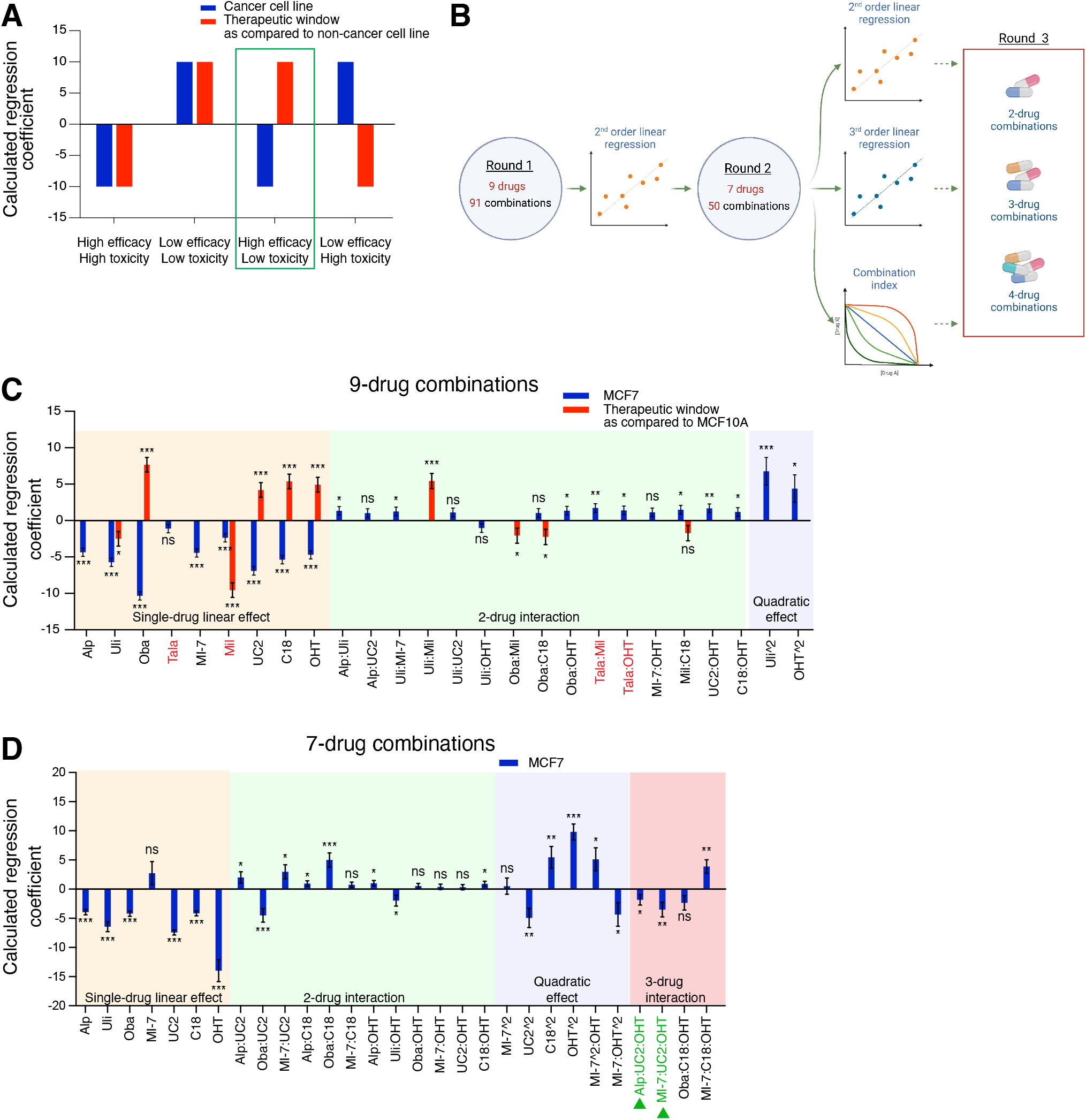
*In vitro* TGMO-based screen identifies synergistic 2-, 3-, and 4-drug combinations for ERα+ breast cancer cells. **(A)** Illustration of the different possible scenarios of the regression modeling of the calculated % ATP levels obtained with cancer cells (blue bars) and the TW compared to a non-cancer cell line (red). An optimal drug or combination of drugs (green frame) would show a negative regression coefficient in cancer cells (high efficacy) and a positive regression coefficient for the TW (low toxicity). Unfavorable drugs or combination of drugs would show either high efficacy and high toxicity, low efficacy and low toxicity, or low efficacy and high toxicity. Illustration was created with GraphPad Prism version 8.0.0. **(B)** Schematic pipeline of the TGMO-based screen. As indicated, the screen was done in 3 rounds. Figure was created with Biorender.com. **(C)** Estimated regression coefficients obtained by second-order linear regression of the calculated % ATP levels from “round 1” with MCF7 cells (blue) and the TW compared to MCF10A cells (red). Drugs highlighted in red in the x-axis labels represent unfavorable profiles of single drugs or of combination of drugs, which were therefore eliminated from the next round. **(D)** Estimated regression coefficients obtained by third-order linear regression of the calculated % ATP levels from “round 2” with MCF7 cells. Drugs highlighted in green in the x-axis labels represent 3-drug combinations with high efficacy in MCF7 cells. For panels C and D, statistical significance is shown as * for *p* ≤ 0.05, ** for *p* ≤ 0.01, *** for *p* ≤ 0.001, and “ns” for non-significant results. Data are represented as means ± standard error of the mean (SEM) of 3 independent experiments, each with 3 independent replicates.

Our pipeline of the combinatorial screen was composed of three experimental rounds (Fig. 2B). From “round 1”, which contains 9 drugs allocated to 91 drug combinations (Fig. S3), second-order regression modeling pointed out drugs that show antagonism, lack of efficacy on cancer cells, and/or toxicity on those that are non-cancerous (Fig. 2C, and Fig. S4A, B). As a result, in the three cell lines MCF7, MCF7-V, and MCF7/LCC2, we eliminated the CDK2 inhibitor milciclib (Bolin *et al*, 2018) from “round 2” since, as a single drug, it showed an antagonistic contribution and an unfavorable TW with MCF10A cells (Fig. 2C, and Fig. S4A, B). The PARP inhibitor talazoparib (Litton *et al*, 2018) showed lack of efficacy as a single drug and/or weak interactions in most of the combinations tested in MCF7 and MCF7-V cells (Fig. 2C, and Fig. S4A). As compared to MCF7/LCC2 cells, the ERK1/2 inhibitor ulixertinib (Germann *et al*, 2017) showed higher toxicity for MCF10A cells (Fig. S4B). In “round 2”, after the elimination of two drugs for each cell line, we continued the combination screen (Fig. 2B) by experimentally testing 50 drug combinations (Fig. S5). The data were analyzed by second-order linear regression (Fig. S6A-C), third-order linear regression, which predicts 2- and 3-drug interactions (Fig. 2D, and Fig. S6D, E), and by calculation of the combination index (CI) based on the Chou Talalay method (Fig. S5), using the CompuSyn software (Chou & Martin, 2005). As a result, our analysis revealed sets of 2-, 3-, and 4-drug combinations that are synergistic in ERα+ breast cancer cell lines and non-toxic to normal cells (Table S2).

### The combination of alpelisib, UC2288, and OHT is synergistic, safe, and selective for ERα+ breast cancer cell lines with activating mutation(s) in PI3Kα

In “round 3”, we tested the 2-, 3-, and 4-drug combinations suggested by “round 2” using a panel of cell lines that correspond to different subtypes of breast cancer (Fig. S7A). We concluded from the dose-response curves of the monotherapies (Fig. S7B-H) that the ERα+/HER2-cell lines were more sensitive to alpelisib, the MDM2 inhibitor MI-773 (Chen *et al*, 2021b), UC2288, and OHT compared to other cell lines (Fig. S7B-E). We analyzed data obtained from “round 3” by calculating the % ATP levels and the CI, and presented the data as heatmaps indicating the differential responses of the different cell lines (Fig. S8). The 2-drug combinations alpelisib + UC2288 and UC2288 + OHT, the 3-drug combinations alpelisib + UC2288 + OHT and talazoparib + UC2288 + OHT, and the 4-drug combination alpelisib + UC2288 + OHT + talazoparib led to reduced ATP levels, increased synergy (based on the calculated CI) and selectivity for ERα+ breast cancer cell lines, including the tamoxifen-resistant MCF7/LCC2 cells, as compared to ERα-negative and non-cancerous cells (Fig. S8C-H). Since low-dose combinations of 3 or 4 drugs might be superior to 2-drug combinations in terms of efficacy and safety, we carefully validated the combination of alpelisib + UC2288 + OHT (AUT), and further evaluated the potential benefit of adding talazoparib to AUT (AUTTala). By measuring % ATP levels we confirmed the efficacy of the AUT drug combination in MCF7 (about 60%), MCF7-V (about 60%), and MCF7/LCC2 (about 65%) cell lines in 72 hour treatments (Fig. 3A-C). Moreover, AUT showed more selectivity for cancer cells than for MCF10A cells (about 15%). Further validation by estimating the cell numbers using crystal violet staining confirmed the efficacy of AUT in MCF7 (about 60%), MCF7-V (about 55%), T47D cells (about 60%), and the two tamoxifen-resistant cell lines MCF7/LCC2 (about 70%) and MCF7/TamR (Knowlden *et al*, 2003) (about 60%) (Fig. 3D-H).

**Fig. 3.**
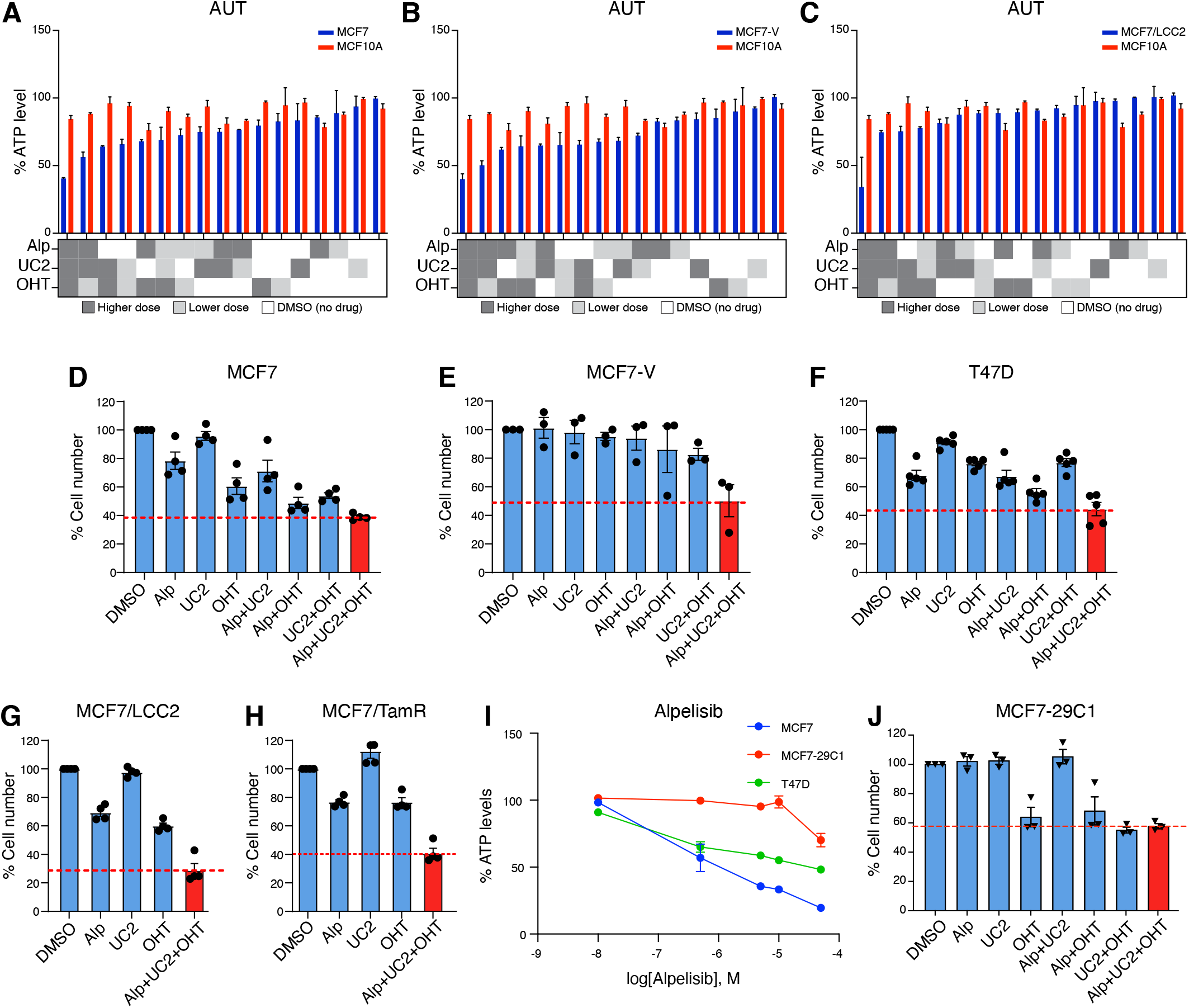
AUT is a synergistic, safe, and selective 3-drug combination for ERα+ breast cancer cells with activating mutation(s) in PI3Kα. **(A-C)** Bar graphs of the % ATP levels of the breast cancer cell lines (blue) MCF7 (panel A), MCF7-V (panel B), and MCF7/LCC2 (panel C) treated with alpelisib (Alp), UC2288 (UC2), and OHT, tested alone or in combinations. Red bars show the results obtained with MCF10A cells. The data are represented as means ± SD of 3 independent experiments. As illustrated in the matrices below the graphs, in addition to the vehicle (DMSO) control in white, each drug was added at two concentrations: higher dose (dark gray), and lower dose (light gray) (Table S1). Values of DMSO only control were set to 100%. In the 3 panels, the same values obtained with MCF10A were used for comparison **(D-H)** Bar graphs of the relative cell numbers for the indicated cell lines treated with alpelisib (Alp), UC2288 (UC2), and OHT, tested alone or in combinations. Bars of the drug combination Alp + UC2 + OHT (AUT) are colored in red. Values of DMSO control were set to 100%. The data are represented as means ± SD (n = 4 independent samples for panel D, G, and H, n = 3 for panel E, n = 5 for panel F). **(I)** Dose-response curves with increasing doses of alpelisib and % ATP levels calculated as the response. The results are shown for MCF7 (PI3Kα-mutant), MCF7-29C1 (wild-type PI3Kα), and T47D (PI3Kα-mutant) cells. For each curve, a control group treated with vehicle was set to 100%. Data are represented as means ± SD of 3 independent experiments. **(J)** Bar graph of experiments with MCF7-29C1 as in panels D-H. (n = 3 independent samples). Drug doses are listed in Table S1.

Activating mutations in PI3Kα occur in approximately 40% of ERα+ breast tumors (Fig. 1D, E) and can be associated with endocrine resistance (Martínez-Saéz *et al*, 2020). Being a selective inhibitor of PI3Kα, alpelisib might affect the selectivity of AUT for PI3Kα-mutant over PI3Kα wild-type breast cancers. To clarify the PI3Kα status for all our breast cancer cell lines, we tested the *PIK3CA* gene for the most common mutations resulting in two amino acid changes in the helical domain (C1616G and G1633A), and another two in the kinase domain (A3140G and A3149T) (Wu *et al*, 2005). We found that all ERα+/HER2-cell lines that we used were PI3Kα-mutants (Table S3). Therefore, we included a variant of the original MCF7 cells (MCF7-29C1) that had previously been corrected to wild-type PI3Kα (Beaver *et al*, 2013). We found that MCF7-29C1 cells showed more resistance to alpelisib as monotherapy and in combinations as compared to MCF7 cells (Fig. 3I, J). Thus, AUT is specific for ERα+ breast cancer with an activating mutation in PI3Kα.

### AUT promotes apoptosis, reduces ERK, NFκB, and HIF1α activities, and is associated with enhanced OHT sensitivity

We explored the mechanisms of cytotoxicity of the drug combination AUT. Using propidium iodide (PI)/annexin V-FITC double staining, we found that AUT treatment for 48 hours increased the apoptotic (annexin V-positive) population of MCF7-V cells as compared to the vehicle control (Fig. 4A, and Fig. S9A). Another subset of cells was positive for PI and negative for annexin V, notably with UC2288 treatment, which might indicate cell death by mechanisms other than apoptosis (Fig. 4A, and Fig. S9A). By immunoblotting, we found an increased activation of caspases 3 and 7, as indicated by reduced levels of their corresponding procaspases, and by an increased level of cleaved caspase 3 (Fig. 4B). To investigate whether alpelisib and UC2288 would enhance the impact of OHT on ERα signaling, we tested the transcriptional activity of ERα using the luciferase reporter plasmid ERE-luc. In MCF7-V and T47D cells, there were no differences between AUT and cells that were treated with OHT alone (Fig. S9B, C). The results were similar with ERα-negative HEK293T cells upon expression of exogenous ERα (Fig. S9D). In contrast, AUT caused a bigger reduction in ERα activity than OHT alone in the tamoxifen-resistant cell lines MCF7/TamR and MCF7/LCC2 cells (Fig. 4C, D).

**Fig. 4.**
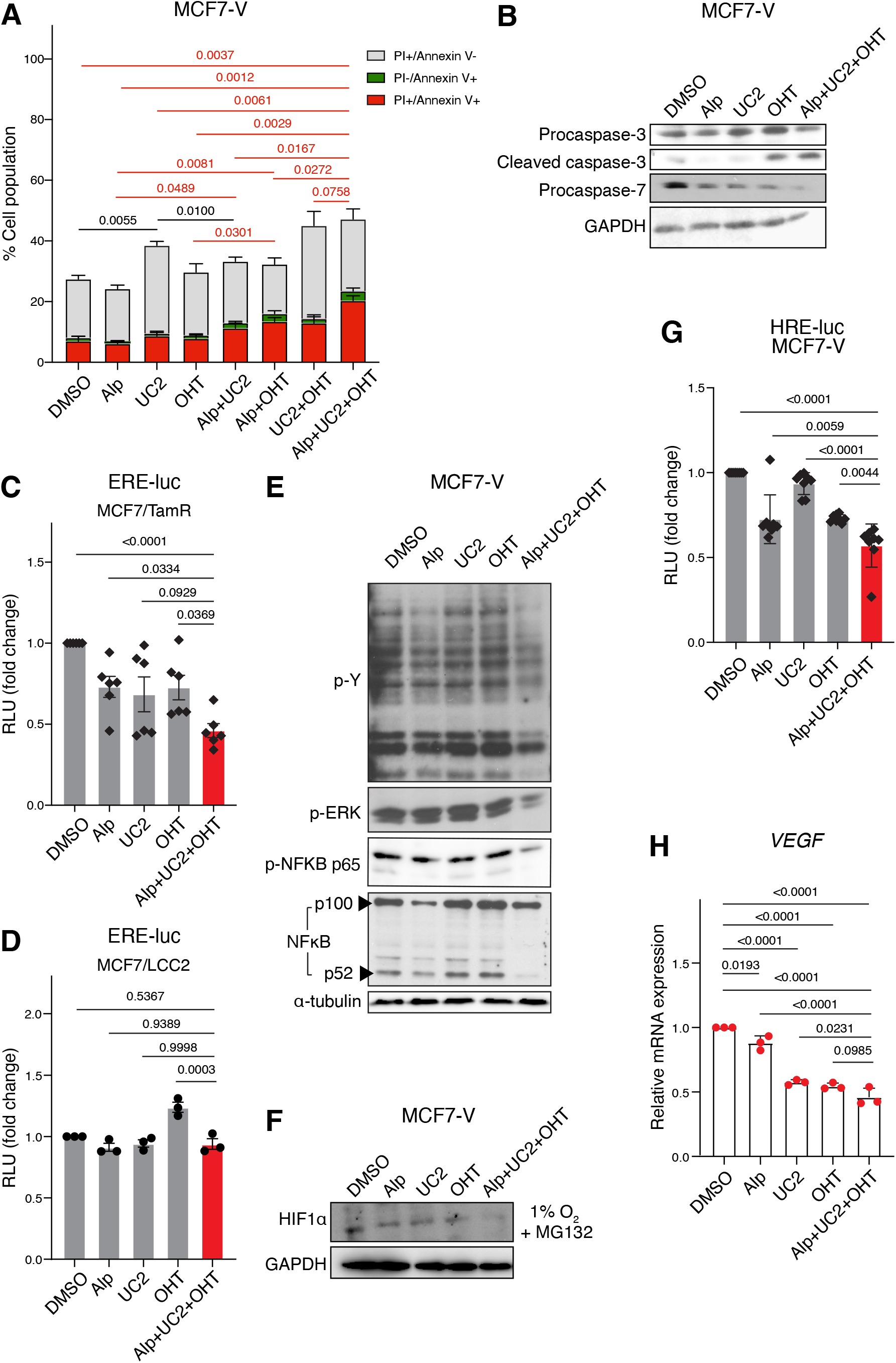
AUT promotes apoptosis, enhances OHT sensitivity, and reduces ERK, NFκB, and HIF1α activities. **(A)** Stacked bar graph of the % cell population of MCF7-V cells differentially stained with PI and annexin V-FITC, as measured by FACS. Cells were treated with alpelisib (Alp), UC2288 (UC2), and OHT, tested alone or in combinations. Total cell population was set to 100%. The data are represented as means ± SD of 3 independent experiments. **(B)** Immunoblots of the indicated proteins in total cell lysates of MCF7-V cells. Cells were treated with alpelisib (Alp), UC2288 (UC2), and OHT, tested alone or in the 3-drug combination. GAPDH was used as an internal standard. **(C,D)** Luciferase reporter assay for the transcriptional activities of endogenous ERα in MCF7/TamR and MCF7/LCC2 cells. Cells were transiently transfected with the ERE-Luc reporter plasmids. Cells were treated with alpelisib (Alp), UC2288 (UC2), and OHT, tested alone or in the 3-drug combination (red bars). Relative luciferase activities (RLU) were normalized to the activities of the internal transfection standard Renilla luciferase. Luciferase activities of the vehicle-treated control groups was set to 1. **(E,F)** Immunoblots of the indicated proteins in total cell lysates of MCF7-V cells. Cells were treated as in panel B. α-tubulin (panel E) or GAPDH (panels F) were used as internal standards. **(G)** Luciferase reporter assay for the transcriptional activities of endogenous HIF1α in MCF7-V cells. Cells were transiently transfected with the HRE-Luc reporter plasmid and treated as in panels C and D. **(H)** *VEGF* mRNA levels measured by RT-qPCR with MCF7-V cells treated as indicated. Ct values were first normalized to *β-ACTIN*, then values corresponding to the DMSO controls were set to 1. For bar graphs, data are represented as means ± SD (n = 3 independent samples for panels D and H, n = 6 for panel C, and n = 8 for panel G). Statistical significance between the groups was analyzed by unpaired student’s t-test for panel A and one-way ANOVA for panels C, D, G, and H. *p*-values ≤ 0.05 were considered statistically significant. Drug doses are listed in Table S1.

Phosphorylation by tyrosine kinases is known to be essential for oncogenic transformation (Kim *et al*, 2017). By immunoblotting for phosphotyrosine residues, we found a reduction in the global phosphorylation of tyrosine residues, indicative of reduced tyrosine kinase activities (Fig. 4E). Since the PI3K/Akt and MEK/ERK pathways are highly interconnected, and their dual targeting was shown to significantly enhance antitumor activity in breast cancer (Ebi *et al*, 2013), we evaluated the effect of AUT treatment on ERK activity. By immunoblotting of phosphorylated ERK, we found that it was synergistically reduced by AUT (Fig. 4E). As a downstream target of PI3K/Akt and MEK/ERK signaling, we investigated the status of the transcription factor NFκB. By immunoblotting, we found that AUT caused a significant downregulation of the active forms of NFκB, indicative of NFκB signaling. We tested for canonical NFκB signaling by immunoblotting for phosphorylated p65. AUT also reduced non-canonical NFκB signaling by reducing the ratio of the active p52 to the inactive p100 NFκB2 family members. Moreover, the phosphorylation of ERK and p65 of NFκB were not induced by the addition of EGF when cells were co-treated with AUT (Fig. S9E). Alpelisib monotherapy seems to be a major contributor to these effects, even though the combination AUT shows a more dramatic synergy. To further validate the effect of AUT on pathways of tamoxifen resistance, we tested the activity of HIF1α under low oxygen conditions (1% O_2_). Immunoblotting of the HIF1α protein, which gets stabilized at low oxygen, showed a reduction in HIF1α induction with AUT treatment, as compared to vehicle as well as to monotherapies (Fig. 4F). Using a hypoxia response element (HRE) luciferase reporter plasmid (HRE-Luc), we found that AUT reduces the maximally induced level displayed by the vehicle control (Fig. 4G, and Fig. S9F, G). Vascular endothelial growth factor (VEGF) is a downstream target gene of HIF1α, which increases angiogenesis and invasiveness of breast cancer cells. Gene expression analysis showed a reduction in VEGF mRNA levels upon single treatment with alpelisib, UC2288, OHT, or with the drug combination AUT (Fig. 4H). Therefore, AUT induces cell apoptosis and reduces major oncogenic signal transduction pathways, such as ERK, NFκB, and HIF1α signaling.

### AUT decreases p21 phosphorylation, induces DNA damage, and impairs cellular senescence

p21 can play dual roles depending on its cellular localization (Parveen *et al*, 2016). In the nucleus, p21 acts as a tumor suppressor (Al-Bitar & Gali-Muhtasib, 2019). However, when Akt phosphorylates p21 at Thr145 and S146, this leads to the stabilization and accumulation of p21 in the cytoplasm (Zhou *et al*, 2001; Rössig *et al*, 2001), where it can bind to and inhibit pro-apoptotic proteins (Suzuki *et al*, 2000). Immunoblotting for p21 phosphorylated on Thr145 revealed a strong reduction with AUT treatment (Fig. 5A). Fluorescent imaging of p21 showed reduced cytoplasmic localization with AUT as compared to vehicle control (Fig. 5B). Most likely, these effects are due to the alpelisib-mediated inhibition of Akt activity.

**Fig. 5.**
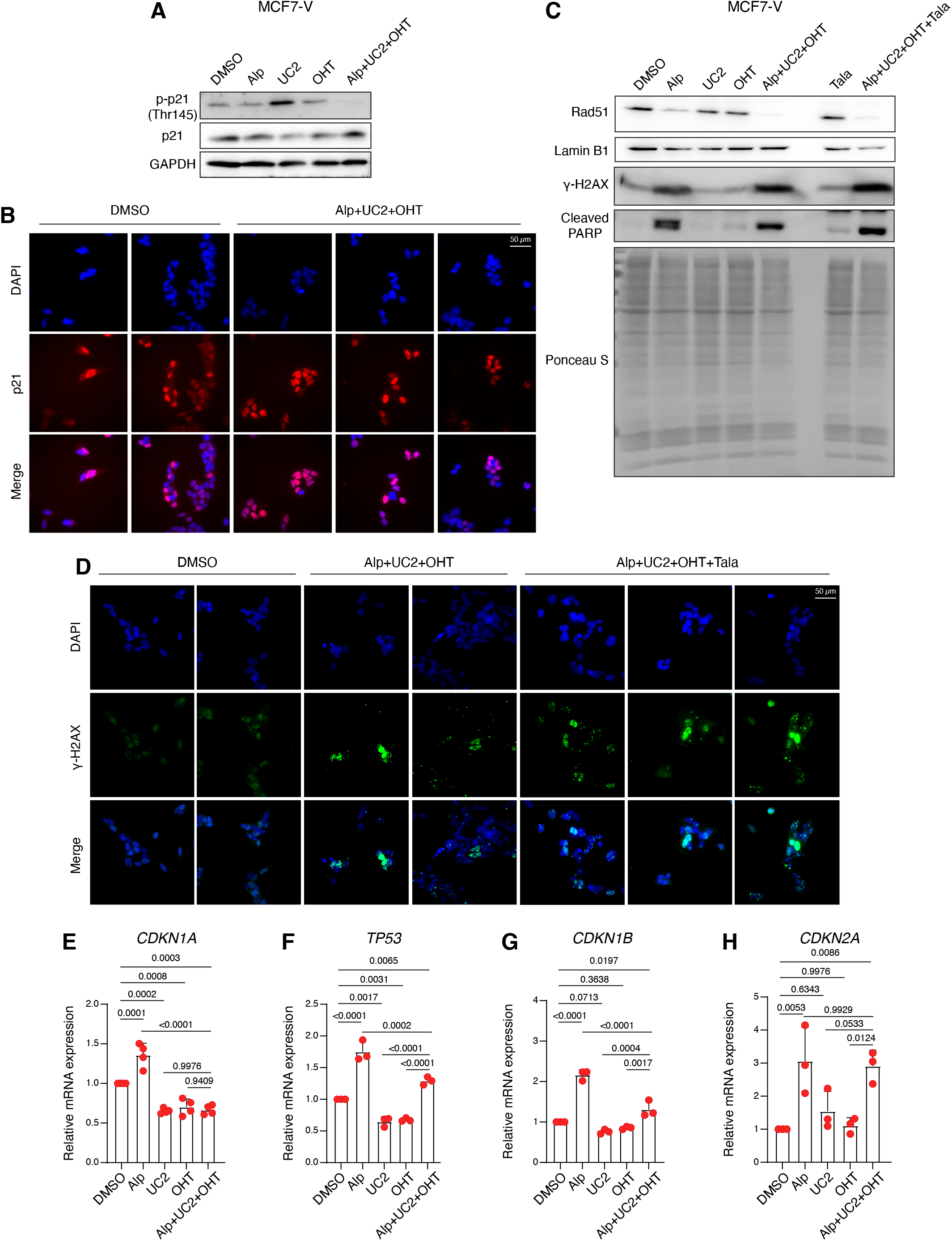
AUT decreases p21 phosphorylation, induces DNA damage, and impairs cellular senescence. **(A)** Immunoblots of the indicated proteins tested in total cell lysates of MCF7-V cells. GAPDH was used as loading control. **(B)** Immunofluorescence images of p21 (red) in MCF7-V cells counterstained for nuclei with DAPI (blue). Cells were treated with DMSO as a control and the 3-drug combination (Alp + UC2 + OHT). **(C)** Immunoblots of the indicated proteins tested in total cell lysates of MCF7-V cells. Cells were treated as in panel A in addition to treatments with talazoaprib (Tala) alone or in the 4-drug combination. Ponceau S-staining of the nitrocellulose membrane was used as loading control. **(D)** Immunofluorescence images of γ-H2AX (green) in MCF7-V cells counterstained for nuclei with DAPI (blue). Cells were treated as in panel B in addition to treatment with the 4-drug combination (Alp + UC2 + OHT + Tala). **(E-H)** mRNA levels of the indicated genes measured by RT-qPCR with MCF7-V cells. Ct values were first normalized to *β-ACTIN*, then values corresponding to the DMSO controls were set to 1. Data are represented as means ± SD (n = 4 independent samples for panel E and n = 3 for panels F-H). Statistical significance between the groups were analyzed by one-way ANOVA and *p*-values ≤0.05 were considered statistically significant. Drug doses are listed in Table S1.

Considering that the inhibition of cytoplasmic p21 may sensitize cells to DNA damage (Pisonero-Vaquero *et al*, 2020), we looked at the DNA damage response. We used the levels of the histone variant *γ*-H2AX as an early marker of DNA double-stranded breaks, PARP1 cleavage as a marker of impaired DNA repair and damage-induced cell death, and the levels of Rad51 as a marker of DNA repair through homologous recombination. We found that aleplisib by itself induces DNA damage and is associated with reduced Rad51 levels and increased cleaved PARP1 levels (Fig. 5C). Treatments with AUT further increased the DNA damage and compromised the repair as indicated by even higher *γ*-H2AX protein levels and lower levels of Rad51, respectively (Fig. 5C, D). Interestingly, the addition of the inhibitor of PARP enzymes (PARP1 and PARP2) talazoparib to AUT further increased the accumulation of DNA damage (Fig. 5C, D). The accumulation of DNA damage leads to cell cycle arrest and eventually to cellular senescence. p21 has been shown to be induced upon DNA damage and to maintain the viability of senescent cells (Yosef *et al*, 2017). We found that alpelisib alone significantly induced the expression of markers of senescence, including *TP53*, *CDKN1A* (p21), *CDKN1B*, and *CDKN2A* (Fig. 5E-H). In contrast, the combination AUT resulted in statistically significantly lower mRNA levels of *TP53*, *CDKN1A*, and *CDKN1B* as compared to alpelisib alone, most probably due to UC2288-mediated inhibition of p21, and hence reduced cell cycle arrest by p53, and reduced senescence. Presumably, the affected cells would eventually die due to the accumulation of DNA damage and impaired DNA repair mechanisms.

### Talazoparib enhances the efficacy of AUT in the long-term treatment of cell lines and in the treatment of patient-derived mammary organoids

To evaluate the potential benefit of adding talazoparib to AUT, we compared the cell numbers after treatment for 72 hours and after long-term treatment for 28 days. We found that talazoparib showed minimal benefit in the 4-drug combination AUTTala in most of the tested cell lines when treated for only a short period (Fig. 6A-E). Long-term treatment of cells with low drug doses can reveal possible benefits in terms of long-term efficacy and evaluate the risk of developing resistance to the drug combination over time. We therefore treated MCF7-V and MCF7/TamR cells with lower doses of each drug in AUT and AUTTala for 28 days. The addition of talazoparib greatly enhanced the efficacy of AUT in the two cell lines (Fig. 6F, G). Therefore, Talazoparib showed a delayed synergistic effect when combined with AUT in cell lines. To explore the efficacy of the drug combinations AUT and AUTTala in a more clinically relevant model, we used *in vitro* patient-derived mammary organoids to mimic as close as possible the tumor microenvironment (Mohan *et al*, 2021; Sachs *et al*, 2018). The organoid model HUB 056 was derived from an ERα+/HER2-/PI3Kα-mutant breast cancer. AUT and AUTTala treatments for one week showed statistically significant reductions of ATP levels of 20% and 30%, respectively, when compared to the vehicle control (Fig. 6H, I). Thus, both drug combinations are efficient in patient-derived organoids, even though even longer periods of treatment can be worth trying in the future.

**Fig. 6.**
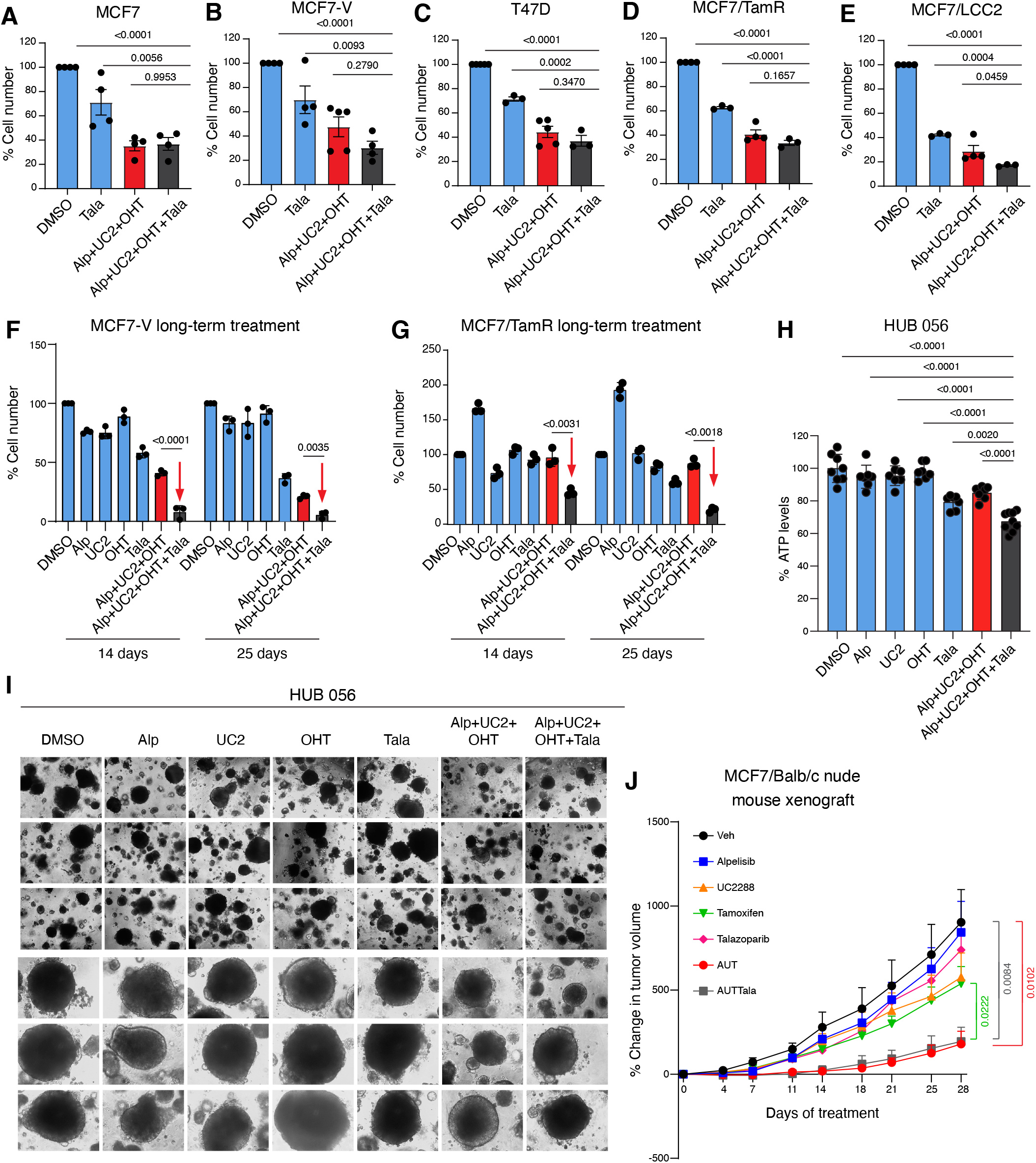
Talazoparib enhances the efficacy of AUT in the long-term treatment of cells and with patient-derived mammary organoids. **(A-E)** Bar graphs of the relative cell numbers of the indicated cell lines treated with talazoparib (Tala) alone or in the 4-drug combination (Alp + UC2 + OHT + Tala) for 72 hours. The 3-drug combination (Alp + UC2 + OHT) was used in parallel for comparison. Values of the DMSO controls were set to 100%. Some values of (Alp + UC2 + OHT) were common with Fig. 3D-H, as they were done in parallel. The data are represented as means ± SD (n = at least 4 independent samples for panels A and B, and n = at least 3 for panels C-E). Statistical significance between the groups were analyzed by one-way ANOVA and *p*-values ≤ 0.05 were considered statistically significant. **(F,G)** Bar graphs of the relative cell numbers of MCF7-V and MCF7/TamR cells treated with alpelisib (Alp), UC2288 (UC2), OHT, and talazoparib (Tala) alone or in the combination of 3 drugs (Alp + UC2 + OHT) or 4 drugs (Alp + UC2 + OHT + Tala). Relative cell numbers measured after 14 days and 25 days of treatment with each drug dose (less than that used for short-term treatments). Values of the DMSO controls were set to 100%. Red arrows point to the results of the efficient 4-drug combination. Data are represented as means ± SD of 3 independent replicates. Statistical significance between the groups were analyzed by two-way ANOVA and *p*-values ≤ 0.05 were considered statistically significant. **(H)** Bar graph of the % ATP levels of patient-derived mammary tumor organoid (model HUB 056) treated with alpelisib (Alp), UC2288 (UC2), OHT, and talazoparib (Tala) alone or in the combination of 3 drugs (Alp + UC2 + OHT) or 4 drugs (Alp + UC2 + OHT + Tala). ATP levels were tested after one week of treatment. The mean value of the DMSO control was set to 100%. Data are represented as means ± SD (n = at least 7 independent samples). Statistical significance between the groups were analyzed by one-way ANOVA and *p*-values ≤ 0.05 were considered statistically significant. **(I)** Representative phase-contrast images of drug-treated organoids after 7 days of treatment. **(J)** Line graphs of the % change in tumor volume of MCF7 xenografts in Balb/c nude mice treated as indicated. Data points represent the means and error bars are SEMs. (n = 5 mice / group). Values on the first day of drug treatments (Day 0) were set to 0%. Differences in the mean values measured on day 28 of treatments were analyzed by unpaired student’s t-test and *p*-values ≤ 0.05 were considered statistically significant. Drug doses are listed in Table S1.

### Activity and safety of AUT and AUTTala in mouse xenografts

To further evaluate the potential benefit of the drug combinations AUT and AUTTala *in vivo*, we set up xenograft experiments with MCF7 cells injected subcutaneously into Balb/c nude female mice. A dose-response study was first performed using two doses for each drug (Fig. S10A, B, and Table S1). Tumor volume was used as a measure of efficacy and the body weight (BW) of animals as an indirect measure of drug toxicity. Except for talazoparib, the selected doses of the drugs were tolerable with a maximum BW loss of 15% with alpelisib at 35 mg/kg (Fig. S10B). In the second phase of the study, we tested the drug combinations AUT and AUTTala. For this, we estimated the individual doses around IC10 to IC20 based on the results of the dose-response experiments (Table S1). After the formation of a palpable tumor, mice were randomized to treatment groups. Treatments with AUT and AUTTala for 28 days showed a statistically significant reduction of tumor growth by about 80%, as compared to vehicle-treated mice, with no significant difference between the two combinations (Fig. 6J). Both drug combinations caused BW loss of about 10% in the treated mice (Fig. S10C). However, demonstrating the benefit of adding talazoparib to AUT, which we had seen *in vitro*, may require variations of the specific treatment regimen. In conclusion, the *in vivo* xenograft model confirmed the efficacy and safety of the drug combination AUT.

## Discussion

Based on our study, we propose two optimized low-dose drug combinations of 3 and 4 drugs, referred to as AUT and AUTTala, respectively, for the treatment of ERα+/HER2-/PI3Kα-mutant breast cancer. PI3Kα mutations occur in about 40% of ERα+ breast cancers, which themselves account for about 70% of all breast cancers. We have identified these drug combinations through sequential screens. We started with an *in silico* screen for potential drug targets using publicly available databases and literature curation, and then used this information to perform *in vitro* combinatorial drug screens using the TGMO-based srceening platform (Weiss *et al*, 2019; Rausch *et al*, 2020a; Nowak-Sliwinska *et al*, 2016). Low-dose synergistic combinations have the advantages of maximizing the efficacy and minimizing the potential toxicity associated with a drug therapy. We have demonstrated *in vitro* that our drug combinations are highly selective and synergistic for ERα+/HER2-/PI3Kα-mutant breast cancer cells, notably compared to non-malignant cells. Most importantly, the drug combinations are also efficient in tamoxifen-resistant breast cancer cell lines, patient-derived mammary tumor organoids, and mouse xenografts with MCF7 cells.

Despite the wide availability of data from cancer patients in public databases, their processing and integration remains challenging (Fillinger *et al*, 2019). To select relevant drug targets in ERα+ breast cancer, we integrated information from multiple datasets that are publicly available (Fig. 1A-E, and Fig. S1A-C). We created a network of SL between genes whose protein products are central in the ERα PPI network (Fig. 1A, B), of genes that are highly co-expressed with ERα (Fig. 1C), and with those that are frequently mutated in ERα+ breast cancer (Fig. 1D, E, and Fig. S1A, B). The synthetic lethal relationships between selected targets in the network make them candidates for efficient and synergistic drug combinations (Fig. 1F and Fig. S1C). We prioritized the drug targets based on their SL score (Fig. 1G), even though the selection for experimental tests was limited by the availability of the relevant inhibitory or stimulatory drugs. Given the complexity of the biological network, the drug targets are interconnected by signaling crosstalk. Collectively these interconnections promote cell growth, proliferation, survival, anti-apoptosis, cell cycle progression, protein synthesis, DNA repair, invasion, metastasis, and angiogenesis (Fig. S1D).

### TGMO-based screen identifies AUT and AUTTala as therapeutically relevant drug combinations

The TGMO-based screen led us to select 4 drugs, 3 of which are FDA-approved for use in breast cancer, that is tamoxifen (Jordan, 2003), alpelisib (Narayan *et al*, 2021) and talazoparib (Hoy, 2018). The fourth drug, UC2288, is in early preclinical studies (Fusco *et al*, 2021; Gupta *et al*, 2014; Wettersten *et al*, 2013). The orally available drug alpelisib is a kinase inhibitor of PI3Kα. UC2288 is a cell-permeable and orally available selective attenuator of p21 expression at the transcriptional or post-transcriptional levels (Wettersten *et al*, 2013). Talazoparib is a potent inhibitor of PARP enzymes which has shown a significant enhancement of progression-free survival in patients with advanced breast cancer with germline *BRCA1/2* mutations as demonstrated by the EMBRACA clinical study (Litton *et al*, 2018). We validated two combinations of the four drugs. Our results demonstrate the efficacy of the combination AUT in reducing the ATP levels (Fig. 3A-C), cell numbers (Fig. 3D-H), and in promoting apoptosis and cell death of ERα+/HER2-/PI3Kα-mutant breast cancer cells (Fig. 4A, B, and Fig. S9A). Moreover, we demonstrate a beneficial role of adding talazoparib to AUT, the combination AUTTala, in long-term treatments (Fig. 6F, G).

Patient-derived organoids resemble organs in structure and function (Sachs *et al*, 2018). We selected a breast tumor model of the ERα+/HER2-/PI3Kα-mutant subtype from a previously established biobank of organoids (Sachs *et al*, 2018). Our results show a significant reduction in the cellular ATP levels of organoids treated with AUT and to a greater extent with AUTTala (Fig. 6H, I). Although the efficacy in organoids was statistically significant, it was lower in magnitude than in cell lines. This could be attributed to the length of the treatment and to the type of assay used to test cell viability. Nevertheless, these results support the notion that our combinations are relevant in 3-dimensional models and effective in the presence of the complex tumor microenvironment. In the future, longer treatment protocols and the detection of more markers, including markers of cell death and growth arrest, could help strengthen the conclusions from this type of model.

Animal models are necessary to recapitulate the pharmacodynamic and pharmacokinetic properties of drugs *in vivo*. Our results demonstrate that the combination AUT with very low doses of the individual drugs is highly efficacious in the reduction of tumor growth of mouse xenografts with MCF7 cells. Moreover, the combination showed high tolerability and safety profiles. With our experimental setting we could not confirm a beneficial role of talazoparib in AUTTala drug combination. This might be due to several factors: (i) The fact that MCF7 cells are highly sensitive to tamoxifen would limit the benefit of talazoparib that was obvious in cases of long-term treatment of tamoxifen-tolerant or -resistant cells (Fig. 6F, G); (ii) the dose and pace of administration of talazoparib may need further optimization since talazoparib showed high toxicity at the selected doses during the initial stages of the study; despite the fact that we further reduced the dose of talazoparib as part of the combination AUTTala by 20x, its dose might still not be optimal; (iii) talazoparib is mainly excreted in urine (Hoy, 2018), which might be incompatible with the use of estradiol pellets, which in the mouse were shown to cause urine retention, obstruction, cystitis, and hydronephrosis (Buhl *et al*, 1985; Walker *et al*, 1992; Pearse *et al*, 2009; Gakhar *et al*, 2009). Although urine retention was prevented by manual bladder expression, hypothetically there could still be some impairment of kidney function, which would lead to increased plasma concentrations of talazoparib, and hence, high toxicity. Based on the available *in vitro* and clinical data for talazoparib, an inhibitory drug-drug interaction among alpelisib, tamoxifen, and talazoparib is less likely, but if anything, it might be potentiating, as will be discussed below. Therefore, further dose optimization, the use of a tamoxifen-resistant model, and assessment of kidney functions might be needed to confirm the advantage of adding talazoparib to AUT *in vivo*.

### Role of PI3Kα and p21 for ERα activity

We show that AUT treatment causes a significant reduction in ERα transcriptional activity when compared to OHT alone in the tamoxifen-resistant cell lines MCF7/TamR and MCF7/LCC2 (Fig. 4C, D). This is not the case in the tamoxifen-sensitive models, which is explicable by the fact that OHT alone already demonstrates a maximal inhibitory effect on the ERα transcriptional activity (Fig. S9B, C). These results are in line with previous studies that highlight roles of PI3K (Campbell *et al*, 2001; Bennesch & Picard, 2015; de Leeuw *et al*, 2011; Alayev *et al*, 2016; Martin *et al*, 2000) and p21 (Fritah *et al*, 2005; Redeuilh *et al*, 2002; Abukhdeir *et al*, 2008) in ERα signaling. As shown by our integrated network of SL (Fig. 1F), in addition to being the most frequently mutated gene, *PIK3CA* appears as one of the genes co-expressed with *ESR1* (ERα), and its inactivating-mutation is synthetically lethal with ERα, which supports a synergistic interaction between their corresponding inhibitors. Alpelisib is the first PI3Kα inhibitor that showed an improvement in the survival of cases with ERα+/HER2-/PI3Kα-mutant advanced or metastatic breast cancers (Narayan *et al*, 2021; Fusco *et al*, 2021). Therefore, it has been approved by the FDA for use in combination with fulvestrant in postmenopausal women with this class of breast cancer (Narayan *et al*, 2021; Bosch *et al*, 2015). Moreover, we found that alpelisib alone and in combinations is more selective for ERα+ breast cancer cells and those with PI3Kα mutation(s) rather than ERα-ones and those with wild-type PI3Kα (Fig. S7B and Fig. 3D-J). This agrees with previous data, which showed that alpelisib preferentially targets mutant PI3Kα (Fusco *et al*, 2021).

Reduced ERα activity can also be explained by referring to the role of p21 in ERα signaling. ERα transcriptional activity mediated by its coactivator CBP was previously shown to be increased by the binding of p21, and that it could be reduced by p21 knockdown (Redeuilh *et al*, 2002; Fritah *et al*, 2005). Another study showed that expression and activity of p21 is necessary for the response to tamoxifen since loss of p21 was associated with tamoxifen resistance, hyperphosphorylation of ERα, and upregulation of ERα target genes (Abukhdeir *et al*, 2008). The relationship between ERα and p21 is bidirectional since p21 expression was shown to be induced by E2, and, not surprisingly, OHT mediates the downregulation of p21 gene transcription (Mandal & Davie, 2010). Therefore, targeting both p21 and ERα could have a great therapeutic benefit.

### Synergistic impact of AUT on signal transduction pathways involved in tamoxifen resistance

Our results highlight a significant impact of AUT on major signal transduction pathways (Fig. S1D) involved in resistance to tamoxifen, such as those involving ERK, NFκB, and HIF1α (Fig. 4E-H, and Fig. S9E-G). PI3K orchestrates the activity of multiple downstream signaling factors. There is crosstalk between PI3K and ERK as indicated by the previously reported downregulation of ERK activity upon inhibition of PI3K (Ebi *et al*, 2013). This agrees with our finding that alpelisib reduces ERK phosphorylation induced by EGF (Fig. S9E). However, the combination of AUT demonstrated a more significant reduction in ERK phosphorylation (Fig. 4E, and Fig. S9E), which highlights the potential of UC2288 in reducing ERK activity even further. This hypothesis is supported by a recent study that showed that UC2288 can inhibit ERK phosphorylation in nasopharyngeal carcinoma (Liang & Zhu, 2021). In addition, AUT displayed a significant reduction in the activity of canonical and non-canonical NFκB signaling (Fig. 4E, and Fig. S9E), which has been shown to greatly impact tamoxifen resistance in breast cancer (Zhou *et al*, 2005; deGraffenried *et al*, 2004; Nakshatri *et al*, 1997). Moreover, we demonstrate that AUT contributes to the downregulation of HIF1α signaling under low O_2_ conditions (Fig. 4F-H, and Fig. S9F, G). It was found that HIF1α contributes to poor prognosis and tamoxifen resistance in breast cancer, and that its inhibition can restore tamoxifen sensitivity (Jögi *et al*, 2019).

### Synergistic targeting of DNA damage response and cellular senescence

We demonstrate that AUT treatment promotes DNA damage (Fig. 5C, D) and is associated with impairment of the damage repair machinery. This is primarily contributed by alpelisib (Fig. 5C). Co-treatment with UC2288 and OHT further augmented DNA damage and reduced repair (Fig. 5C). The effects of alpelisib on the DNA damage response are most likely due to the inhibition of Akt activity downstream of PI3Kα inhibition. There are accumulating data supporting the role of Akt activity in the DNA damage response. Akt activity is correlated with resistance to genotoxic agents and ionizing radiation in multiple types of cancer due to its positive role in DNA damage repair (Kao *et al*, 2007; Toulany *et al*, 2008; Li *et al*, 2009; Liu *et al*, 2014). Upon DNA damage, sensors, such as ATM, ATR, DNA-PK, and PARP1, cause activation of Akt, which further leads to cell cycle arrest, inhibition of apoptosis, and DNA repair (Liu *et al*, 2014). Since DNA damage induces p53 activity which induces p21, p21 inhibition by UC2288 is very relevant. UC2288 on its own was recently shown to promote DNA damage in nasopharyngeal carcinoma (Liang & Zhu, 2021). Induction of p21 allows the repair of the damage by halting cell growth and inhibiting apoptosis simultaneously. Reduction of p21 levels sensitizes cancer cells to DNA-damaging agents, such as doxorubicin, etoposide, camptothecin, and *γ*-irradiation. It prevents DNA damage-induced G1 arrest, causes an initial blockage of the cell cycle in G2 followed by rounds of DNA synthesis without mitosis, and finally leads to apoptosis (Waldman *et al*, 1996; Waldman *et al*, 1997; Tian *et al*, 2000). Attenuation of the cytoplasmic activity of p21 can also be achieved indirectly by the inhibition of Akt with alpelisib in the AUT combination. In Akt knockout mice the induction of p21 due to DNA damage is impaired (Bozulic *et al*, 2008). Thus, co-targeting of Akt and p21 can strongly impair DNA damage repair (Fig. S1D).

DNA damage contributes to cellular senescence, which can stabilize cancer rather than induce its regression (Te Poele *et al*, 2002; Hanahan, 2022). p53 and p21 play important roles in the induction of senescence (Te Poele *et al*, 2002). p21 levels were found to be directly correlated with cellular senescence in HeLa cells, where p21 downregulation was associated with decreased senescence (Wells *et al*, 2000; Goodwin & DiMaio, 2000) and re-entry of cells into S-phase, but not cell division (Ma *et al*, 1999). Cleavage of p21 by caspase-3 in response to DNA damage was shown to shift the cell growth arrest to apoptosis (Zhang *et al*, 1999). Our results show that alpelisib induces the mRNA levels of the markers of senescence, whereas the combination of AUT reduced this induction (Fig. 5E-H). Inhibition of p21 by UC2288 as part of the AUT combination would lead to decreased cellular senescence induced by DNA damage. Therefore, impaired DNA damage repair, impaired senescence, and accumulation of DNA lesions will eventually lead to apoptosis (Fig. 4A, B), most likely to preserve the genomic integrity of the remaining cell population (Weiss, 2003).

### Advantages of adding talazoparib to the AUT drug combination

The results of the TGMO-based screen pointed out that talazoparib can have a higher impact in the tamoxifen-resistant setting (Fig. 2C, and Fig. S4A, B). By further investigations, we identified a great benefit of adding talazoparib to AUT when given at a low dose and for a longer treatment period in MCF7-V and MCF7/TamR cells (Fig. 6F, G). This can be partly attributed to the known role of PARP1 in ERα signaling (Schiewer & Knudsen, 2014; Zhang *et al*, 2013; Pulliam *et al*, 2019; Gadad *et al*, 2021). PARP1 plays a role in chromatin remodeling and regulation of gene transcription in breast cancer (Gibson *et al*, 2016; Krishnakumar *et al*, 2008; Frizzell *et al*, 2009; Krishnakumar & Kraus, 2010). It is required for the transcriptional activity of ERα, promotes the transcription elongation of estrogen-regulated genes by RNA polymerase II, and its absence would reduce the binding of ERα and FOXA1 to ERα binding sites (Gadad *et al*, 2021; Ju *et al*, 2006; Schiewer & Knudsen, 2014). ERα was also shown to be poly-ADP-ribosylated by PARP1 within the DNA binding domain in vascular smooth muscle cells, which enhances the nuclear localization of ERα (Zhang *et al*, 2013). The idea of co-targeting of ERα and PARP1 is supported by reports that showed that a combination of tamoxifen and talazoparib sensitized breast cancer cells to tamoxifen through the reduction of ERα poly-ADP-ribosylation (Pulliam *et al*, 2019).

In addition to the impact of PARP1 inhibition on ERα activity, talazoparib leads to PARP1 being trapped at sites of DNA damage. In our SL network, the *PARP1* and *BRCA1* genes appear as the top synthetic lethal gene pair (Fig. 1F, G). Inhibition of PARP1 in *BRCA1/2* mutant breast cancer was shown to induce cell death due to the accumulation of irreparable DNA damage. PARP1 activity was found to be increased in tamoxifen-resistant cells due to the increased levels of reactive oxygen species (ROS) (Pulliam *et al*, 2019). Tamoxifen treatment was associated with ROS-mediated impairment of homologous recombination repair, upregulation of PARP1 levels and accumulation of DNA damage (Qi *et al*, 2017; Pulliam *et al*, 2019). This further strengthens the potential benefit of a combination of PARP1 inhibitor and tamoxifen. In line with this, our results show that the addition of talazoparib to AUT maximized the accumulation of DNA damage seen with AUT (Fig. 5C, Fig. S1D). The delayed impact of talazoparib in the combination AUTTala could be explained by the time needed for the accumulation of DNA damage and the heterogeneity of the responses of individual cells.

### Potential for pharmacokinetic drug-drug interactions

Apart from the above-mentioned pharmacodynamic interactions, drug interactions could also happen at the level of their pharmacokinetics. Modulation of drug absorption, distribution, metabolism, and/or excretion can affect the bioavailability of the active drug, and hence its efficacy and/or toxicity. In AUT and AUTTala, since 3 drugs are approved by the FDA, some information is available to help predict pharmacokinetic interactions. Tamoxifen is highly metabolized by liver enzymes, including the cytochrome P450s CYP2C9, CYP2D6, and CYP3A4, to its 30-100x more active metabolites. The major metabolic pathways of tamoxifen are the N-de-methylation (around 92%) and 4-hydroxylation (around 7%), which yield N-desmethyltamoxifen and OHT as primary active metabolites, respectively (Klein *et al*, 2013; Lien *et al*, 1989). These metabolites undergo a second conversion to another highly potent form, 4-hydroxy-N-desmethyltamoxifen (known as endoxifen), which represents the most abundant metabolite (Klein *et al*, 2013; Lien *et al*, 1989).

Alpelisib was shown with *in vitro* assays to induce CYP2C9 and CYP3A4 (European Medicines Agency (CHMP), 2020), which are involved in the conversion of tamoxifen to OHT and N-desmethyltamoxifen. A minor proportion of alpelisib is metabolized by the enzyme CYP3A4 (James *et al*, 2015), which can be affected by tamoxifen co-administration. Since tamoxifen can inhibit the activity of this enzyme, the alpelisib concentration might be raised. Moreover, tamoxifen, alpelisib, and talazoparib are substrates of the efflux pumps p-glycoprotein (Callaghan & Higgins, 1995; European Medicines Agency (CHMP), 2020; Elmeliegy *et al*, 2020) and BCRP (European Medicines Agency (CHMP), 2020; Guney Eskiler *et al*, 2020; Selever *et al*, 2011) at varying affinities, which might allow competitive interactions. Efflux pumps are widely present in the cells of multiple organs, such as the small intestines, tubules of the kidneys, and the liver, in addition to the cancer cells themselves, which can render them drug resistant. Inhibition of efflux pumps by one or more of the drugs might increase the plasma concentrations of the others and/or itself and increase the intracellular drug concentration in cancer cells. Therefore, monitoring of therapeutic doses and plasma concentrations might be recommended.

Taken together, the drug combinations proposed in this study might be of great therapeutic value and potential alternatives to current approaches for ERα+ breast cancer. Drug-associated toxicity and the development of resistance are major obstacles to efficient cancer therapy. For example, the FDA-approved combinations of fulvestrant + CDK4/6 inhibitors can be associated with high grade hematologic toxicities such as neutropenia and leukopenia (Lorfida *et al*, 2020). The emergence of some driver gene mutations during treatments was detected by circulating tumor DNA analysis, revealing mutations in *RB1* and to a larger extent in *PIK3CA* and the ERα gene *ESR1* itself (O’Leary *et al*, 2018a; O’Leary *et al*, 2018b). Moreover, high levels of cyclin E1 mRNA (*CCNE1*) were associated with poor response to palbociclib (Turner *et al*, 2019). In the combinations of antiestrogens + alpelisib, high grade adverse events, including hyperglycemia, rash, and diarrhea are common and mutations in *PTEN* and *ESR1* were associated with disease progression and resistance to the combinations (Rugo *et al*, 2020; Razavi *et al*, 2020). A preclinical study revealed a great benefit of adding alpelisib to overcome resistance to the dual therapy of fulvestrant + CDK4/6 inhibitors (O’Brien *et al*, 2020). A similar combination of the PI3K inhibitor taselisib with fulvestrant and palbociclib showed greater efficacy than fulvestrant + palbociclib in a phase I clinical trial (Pascual *et al*, 2021). More and more, higher order combinations are seen as more promising than dual therapies. Based on our study, we propose the co-targeting of ERα, PI3Kα, p21, and PARP1 activities as a potentially efficient and safe strategy for blocking multiple cell survival avenues, reducing the emergence of resistance, and eventually leading to the selective death of ERα+ breast cancer cells.

## Materials and methods

### Construction and analysis of ERα PPI network

Using the software Cytoscape 2.6.3 (Han *et al*, 2004; Shannon *et al*, 2003), the list of 322 interactors of ERα (see updated list at http://www.picard.ch/downloads was given a feature named “DP” (Data S2) and was imported into the human interactome network that was previously developed by our lab (Echeverría *et al*, 2011). The two lists were merged by union as follows: Tools-> merge-> Network. A new network that contains all interactors with ERα and all known interactions between them was extracted as follows: selecting the feature DP and then: File -> New -> Network -> From selected nodes, all edges. The new network represents the ERα interactome. The layout of the network was arranged based on weighed edges as follows: Layout -> Edge Weighted Spring Embedded Layout (Data S1). For the analysis of the network, all interactors were selected and then analyzed for node centrality measures as follows: Tools -> Network Analyzer -> Network Analysis -> Analyze Network -> Treat the Network as undirected. Analysis was exported as excel file (Data S3) and several parameters were investigated such as number of directed edges, average shortest path, radiality and stress. Based on these parameters, the list was ranked and frequency distribution histograms were constructed (Fig. 1B). The top 50 interactors were selected from the network and highlighted in green (Fig. 1A and Data S1).

### Mining gene expression and mutational profiles in ERα+ breast cancer

Genes that are co-expressed with ERα were mined using the online databases Oncomine (Rhodes *et al*, 2004) and GOBO (Ringnér *et al*, 2011). The list of genes was ranked based on the co-expression coefficient. The mutational landscape of ERα+ breast cancer was mined using the publicly available Catalogue Of Somatic Mutations In Cancer (COSMIC) (Ringnér *et al*, 2011) and Integrative Onco-Genomics (IntOGen) (Perez-Llamas *et al*, 2011) databases.

### Construction of SL network

Synthetic lethal relationships in breast cancer from the two databases SynLethDB (Guo *et al*, 2016) and the siRNA-validated list of cancer-driving synthetic lethal gene pairs (Ye *et al*, 2016) were merged with Microsoft Excel, and a SL network (Fig. S1C) was constructed by importing the list to Cytoscape. In the network, genes that correspond to ERα interactors, are co-expressed with ERα, and genes that are frequently mutated were highlighted and given a color code (Fig. S1C and Data S5). For a more focused search for drug targets, we eliminated those genes that were not identified as ERα interactors, as being co-expressed, or mutated in ERα+ breast cancer. Therefore, a smaller SL network was constructed (Fig. 1F and Data S5). In this network, genes that happened to be at the top of any of the lists of ERα interactors, of genes co-expressed with ERα, or of mutated genes were further labelled with a bold red outline (Fig. 1F).

### Selection of drugs

Screening for drugs was done using the literature and databases, including DRUGBANK (Wishart *et al*, 2008), ChEMBL (Mendez *et al*, 2019), Guide to PHARMACOLOGY (Harding *et al*, 2022). Details about the drugs and the doses used in the screen or in other experiments are listed in Table S1. Stock concentrations were adapted so that the maximum DMSO concentration in a combination is ≤ 0.1%. Aliquots of 20 μl were stored at −20°C and thawed directly before use.

### Cell culture and cell lines

The following cell lines were from ATCC: MCF7 (HTB-22), BT474 (HTB-20), SKBR3 (HTB-30), MDA-MB-231 (HTB-26), HCC1937 (CRL-2336) (Foray *et al*, 2002), MCF10A (CRL-10317) (a gift from Eric Allémann, University of Geneva), HEK 293T cells (CRL-3216), RPE1 cells (CRL-4000), and CCD-18Co (CRL-1459). T47D cells were purchased from Sigma-Aldrich (#85102201). MCF7-V cells, which are related to wild-type MCF7 cells, notably differ from them by being more OHT-tolerant and displaying more robust ERα responses (Mohammadi Ghahhari *et al*, 2022; Hany *et al*, 2022). We used MCF7/LCC2 (Brünner *et al*, 1993), and MCF7/TamR cells (Knowlden *et al*, 2003) (a gift from Denise Barrow, Cardiff University) as tamoxifen-resistant cell lines. H3396 cells (Garrigues *et al*, 1993) were a gift from Hinrich Gronemeyer’s laboratory (IGBMC, Strasbourg). MCF7-29C1 (Beaver *et al*, 2013) were a gift from Ben Ho Park of Vanderbilt University Medical Center.

The tamoxifen-resistant cell lines MCF7/LCC2 and MCF7/TamR were cultured in hormone-deprived medium containing 100 nM OHT (Sigma-Aldrich #H7904). MCF10A were cultured in DMEM/F12 medium (Thermo Fisher Scientific # 31331028) containing 5% horse serum (Bio-Concept #2-05F00-I), 100 μg/ml penicillin/streptomycin, 10 ng/ml human epidermal growth factor (Sigma-Aldrich #E9644), 5 μg/ml insulin (Sigma-Aldrich #I9278), and 1 μM dexamethasone (Sigma-Aldrich #D8893). The rest of the cell lines were cultured in DMEM/GlutaMAX medium (Thermo Fisher Scientific #31966047), containing 10% fetal bovine serum (PAN-Biotech #P40-37500), and 100 μg/ml penicillin/streptomycin. All cell lines were maintained at 37°C with 5% CO_2_ in a humidified incubator.

### Cell viability assay

Cells were hormone-starved 72 hours prior to treatments by incubation in “hormone-deprived” medium made of phenol red-free Dulbecco’s Modified Eagle’s Medium (DMEM) (Thermo Fisher Scientific #11880036), containing 5% charcoal-treated fetal bovine serum (CHFBS), 100 μg/ml penicillin/streptomycin (Thermo Fisher Scientific #15070063), 4 mM L-glutamine (Thermo Fisher Scientific #25030081). One day prior to treatments cells were seeded into 96-well plates at the following cell numbers/well: 12,000 for MCF7, 15,000 for MCF7-V and for MCF7/LCC2, 6,000 for MCF10A, 4,000 for CCD-18Co and for RPE1. For the dose-response assays, cells were treated in triplicates with a serial dilution of each drug prepared in hormone-deprived medium containing 5 pM E2. For the combinatorial assays, in addition to the vehicle (DMSO) control, each drug was given at 2 dose levels (Table S1). After 72 hours, ATP levels were measured with the CellTiter-Glo (CTG) reagent (Promega #G7572) according to the manufacturer’s instructions. Cell lysates were transferred to black opaque 96-well plates (Fisher scientific #50-905-1574), and luminescence was measured using a Cytation 3 Image Reader (Agilent). The luminescence of vehicle-treated controls was set to 100%. Data were fitted by least squares regression with a variable slope (four parameters) using GraphPad Prism version 8.0.0. Calculation of IC20 was done by interpolation of the dose-response curves with a 95% confidence interval. TW was calculated as follows: (TW = %ATP in non-cancerous cells (MCF10A) - %ATP of cancerous cells).

### TGMO-based screen

As previously described (Weiss *et al*, 2019), cells were treated for 72 hours with the corresponding treatments using a pre-designed matrix. Cell viability was inferred from the measurement of the ATP levels done as described above. MCF10A cells were treated in parallel to help calculate the TW.

### Regression analysis in the TGMO-based screen

%ATP levels were modelled by second-order or third-order linear regression using the software MATLAB. The calculated regression coefficients quantify the contribution of each drug alone (single-drug linear effect and single-drug quadratic effect) and the overall contributions of 2-drug combinations (2-drug interaction), or 3-drug combinations (3-drug interaction in case of a third-order linear regression analysis). Similarly, we calculated the regression coefficient of the TW (Fig. 2A). Negative coefficients indicate a synergistic contribution, while positive coefficients indicate an antagonistic contribution of the drug or its combinations (Fig. 2A). Therefore, an ideal drug combination would have a negative regression coefficient in cancer cells and a positive regression coefficient for the TW (Fig. 2A). The predictive strength of the models was determined by multiple statistical measures, such as the correlation coefficient R^2^, residual analysis, normal distribution, and analysis of outliers by Cook’s distance (Nowak-Sliwinska *et al*, 2016). Statistical significance of the calculated regression coefficients was calculated by ANOVA lack-of-fit test and *p*-values ≤ 0.05 were considered statistically significant.

### Calculation of CI

In the TGMO-based screen, the fraction affected (Fa) by single-drug or combination treatments was calculated by the following equation: 1 - (luminescence of drug-treated wells / luminescence of vehicle-treated wells). Using the software CompuSyn, Fa values for each drug combination were entered to calculate CI values.

### Mammary tumor organoids

The mammary tumor organoid model HUB 056 was purchased from HUB organoids (HUB, the Netherlands). The protocol was adapted from the provider’s protocol (Sachs *et al*, 2018). Briefly, organoids were resuspended in a mixture of 80% Cultrex Basement Membrane Extracts (BME) (Amsbio/Trevigen, # 3533-010-02) and 20% mammary tumor organoid medium (MTM). MTM used to maintain organoid culture is composed of Ad-DF+++ medium (Advanced DMEM/F12 (ThermoFisher, # 12634028) (Ad-DF) supplemented with 4 mM L-glutamine, 100 μg/ml penicillin/streptomycin, and 10 mM HEPES buffer pH 7.4), 1.25 mM N-acetyl cysteine (Sigma Aldrich, #A9165), 500 nM of the TGFβ receptor inhibitor A83-01 (Tocris, #2939), 1x B27 supplement (ThermoFisher, #17504044), 5 ng/ml human EGF (Sigma Aldrich, #E9644), 5 ng/ml human KGF/FGF-7 (Peprotech, #100-19), 20 ng/ml human FGF-10 (Peprotech, #100-26), 37.5 ng/ml human heregulin-β 1 (Peprotech, #100-03), 2% noggin-conditioned medium (harvested using the HEK 293T/Noggin cell line obtained from the Hubrecht institute, the Netherlands), 10 mM nicotinamide (Sigma Aldrich, #N0636), 50 μg/ml Primocin (InvivoGen, #ANT-PM-2), 250 ng/ml recombinant human R-Spondin 3 (Peprotech, #120-44), 500 nM of the p38 inhibitor SB 202190 monohydrochloride hydrate (Peprotech, #1523072), 5 μM Rho kinase inhibitor Y-27632 dihydrochloride (Peprotech, #1293823). The suspension of organoids in BME/Ad-DF+++ was seeded in the form of domes (i.e. drops of 10 μl each) into 24-well plates, which were pre-warmed overnight at 37°C. Plates were kept inverted at 37°C for 30 min or until the domes solidified, then 500 μl MTM / well were carefully added and plates were maintained at 37°C with 5% CO_2_ in a humidified incubator. Organoids were passaged (split ratio 1:2) by mechanical shearing approximately every 20 days or when almost 80% of organoids were >50 μm. Organoids were visualized using an inverted light microscope and images were captured using a Dino-lite camera and the software DinoXcope.

### Cell viability assay with organoids

Cell viability was determined by measuring ATP levels using the CellTiter-Glo 3D Cell Viability Assay (Promega, #G9683). Briefly, domes of organoids/BME suspension (5 μl) were seeded as one dome per well of a pre-warmed 96-well plate. After adding 60 μl MTM to the organoids, they were maintained at 37°C with 5% CO_2_ in a humidified incubator. After 2 days of seeding, drugs were prepared in 2x concentrated working solutions and then added to the wells in equal volume to make the final concentration of the drugs 1x. After one week of drug incubation (Table S1), the CellTiter-Glo 3D reagent was added, and the plates were vigorously shaken for 30 min while being protected from light. Lysates were transferred to black opaque 96-well plates and luminescence was measured using a Cytation 3 Image Reader. The luminescence of vehicle-treated control was set to 100%.

### Crystal violet assay

Cells were seeded into 12-well plates and treated for 72 hours with the corresponding treatments (Table S1). For long-term treatments, cells were seeded into 6-well plates and split once per week at a ratio of 1:3. Relative cell numbers were assessed using crystal violet. Adherent cells were washed twice with phosphate-buffered saline (PBS) and fixed with 4% formaldehyde in PBS. After 20 min, cells were washed twice with PBS and then 0.1% crystal violet solution in distilled water was added and left on the cells for 30 min. Crystal violet was carefully discarded. Stained cells were washed with distilled water and left to air-dry overnight. Glacial acetic acid was added to dissolve the crystals and the absorbance was measured at 595 nm with a Cytation 3 Image Reader (Agilent). The values of vehicle-treated controls were set to 100%.

### PI / annexin V staining and flow cytometry

Cells were seeded at a density of 0.5 x 10^6^ cells / well. After treatments for 48 hours (Table S1), cells were trypsinized, washed with PBS, and resuspended in 100 μl annexin V-binding buffer that contains 10 mM HEPES pH 7.4, 150 mM NaCl, and 2.5 mM CaCl_2_. Then, 5 μl annexin V-FITC and 2.5 μg/ml PI were added to the cells and kept for 20 min at 4 °C. Cells were analyzed by flow cytometry using a FACS Gallios (Beckman Coulter). Data were analyzed using ths software FlowJo. The distribution of the cell population was divided by gating into 4 quadrants. One quadrant contains unstained healthy cells, another quadrant cells only stained with annexin V-FITC corresponding to early apoptotic cells, another quadrant contains cells stained both with annexin V-FITC and PI corresponding to late apoptotic cells, and the fourth quadrant contains cells only stained with PI, which indicates necrotic cell death or death by mechanisms other than apoptosis.

### Immunoblot analysis

For assays of HIF1α, 18 hours after drugs had been added, cells were further treated with 1 μM of the proteasome inhibitor MG132 (UBPBio, # F1101), and then maintained in only 1% O_2_ for 6 hours. For assays with EGF, cells were treated with the indicated drugs for 24 hours and EGF (10 ng/ml) was added 15 min before harvesting the cells to induce MAPK signaling. For the rest of the assays, cells were treated as indicated for 24 hours at 37°C, 5% CO_2_ (Table S1). Cell pellets were then harvested and washed twice with PBS. Cells were lysed in ice-cold lysis buffer, which contained 20 mM Tris-HCl pH 7.4, 2 mM EDTA, 150 mM NaCl, 1.2% sodium deoxycholate, 1.2% Triton-X-100, protease inhibitor cocktail (Thermo Fisher Scientific #78429), and phosphatase inhibitor cocktail (Thermo Fisher Scientific #78420)). Lysates were sonicated at high power with a Bioruptor sonicator (Diagenode) for 15 min. The concentration of total proteins was determined using the Bradford reagent (Biorad #5000001) and measuring absorbance at 595 nm. Using SDS-polyacrylamide gel electrophoresis, a volume corresponding to 20-50 μg of proteins was loaded on the gels and proteins resolved by electrophoresis. Then, proteins were transferred from the gel to a nitrocellulose membrane (GVS Life Science) using an electroblotting unit (VWR #BTV100). Blocking was done for 20-60 min in 2.5% non-fat dry milk in Tris-buffered saline with 0.2%Tween-20 (TBST) or using bovine serum albumin (BSA) in TBST in the case of phosphorylated proteins. Primary antibodies were added to the membranes and kept overnight at 4°C. Membranes were washed at least 3 times with TBST for 1 hour. Secondary antibodies were added at room temperature for 1 hour. Protein bands were visualized using the WesternBright^TM^ chemiluminescent substrate (Advansta #K-12045-D50) and X-ray films or using an Amersham™ ImageQuant™ 800 biomolecular imager. Molecular weights were determined using the “Fisher BioReagents EZ-Run Prestained Rec Protein Ladder” (Thermo Fisher Scientific). A list of primary and secondary antibodies is available in Table S4.

### RNA extraction and real-time RT-qPCR

RNA was extracted using a homemade guanidinium-acid-phenol reagent (Chomczynski & Sacchi, 1987). Cells were seeded at a density of 0.5 x 10^6^ cells / well. After drug treatments (Table S1), cells were harvested and lysed in 500 μl solution D (4 M guanidium isothiocyanate, 25 mM sodium citrate pH 7 and 0.5% N-laurosylsarcosine, and 0.1 M β-mercaptoethanol). Then, for each tube, 50 μl of 2 M sodium acetate pH 7.4, 500 μl water-saturated phenol (ROTH #A980.1), and 100 μl chloroform/isoamyl alcohol (49:1) (Merck, #25668) were added. Tubes were shaken vigorously and then centrifuged for 20 min at 10,000 g, 4°C. The top aqueous phases were recovered into new microfuge tubes and RNA was precipitated by adding 250 μl isopropanol followed by centrifugation for 30 min at 12,000 g, 4°C. Pellets of RNA were washed with 250 μl of 75% ethanol. Reverse transcription of RNA to cDNA was done by mixing 400 ng total RNA, random primers (Promega #C1181), and GoScript reverse transcriptase kit (Promega #A5003). Then, GoTaq qPCR Master Mix kit (Promega #A6001) and specific primers were used to set up the quantitative real-time PCR (qPCR) reactions according to the manufacturer’s protocol. We used a Bio-Rad CFX 96 Real-Time PCR instrument (Bio-Rad) and calculated the Ct values. Relative gene expression was calculated using the ΔΔCt method. A list of the specific primer sequences is in Table S5.

### Dual luciferase reporter assay

Cells were seeded at 60% confluency in 6-well plates and incubated in hormone-deprived medium for 72 hours. Using the transfection reagent PEI MAX 40K (Polysciences Inc. # 24765-100), cells were co-transfected with 50 ng of the Renilla luciferase transfection control plasmid pRL-CMV (Promega #E2261) and 1 μg of either the ERα reporter plasmid XETL (here referred to as ERE-Luc) (Bunone *et al*, 1996) or the HIF1α reporter plasmid HRE-Luc (Emerling *et al*, 2008) (a gift from Navdeep Chandel (Addgene #26731)). 24 hours later, cells were harvested and seeded into 96-well plates. 24 hours after seeding, cells were treated with the corresponding drugs for another 24 hours (Table S1). For assays using HRE-Luc, cells were incubated in 1% O_2_ using a hypoxia incubator during the drug treatments. For assays in HEK 293T cells, 1 μg of the plasmid HEG0 (Tora *et al*, 1989), which expresses full-length ERα, was added to the PEI MAX transfection mixture mentioned above. In parallel, a group of cells were transfected with the plasmid pSG5 (Green *et al*, 1988) as the empty vector control. Using the dual-luciferase detection kit (Promega #E1910), cells were lysed in the lysis buffer for 30 min, and luciferase activities were measured with Cytation 3 Image Reader (Agilent). Firefly luciferase activities were normalized to those of the Renilla luciferase.

### Immunofluorescence and imaging

Cells were seeded onto sterile glass coverslips in 12-well plates at a density of 0.2 x 10^6^ cells per well. After 24 hours of drug treatments (Table S1), cells were washed twice with PBS and then fixed with 4% formaldehyde for 20 min at room temperature. After one wash with PBS, the coverslips were washed twice with TBS. For permeabilization, cells were kept in 0.1% Tritone-X100 in TBS for 4 min at room temperature. Cells were washed once with TBS and then blocked with 1% BSA in TBS for 1 hour. Primary antibodies were diluted in 1% BSA in TBS and added to the cells overnight, at 4°C, in a humidified chamber. Cells were washed in TBS with gentle agitation for 10 min. Fluorescent secondary antibodies were diluted in 1% BSA in TBS and added to the cells for 20 min to 1 hour at room temperature, in a dark chamber. Cells were washed with TBS for 10 min and then then nuclei were counterstained by incubation with diamidino-2-phenylindole dye (DAPI) (1:30,000 in PBS, from a 1 mg/ml stock solution, Thermo Fisher Scientific #62248) for 5 min. Cells were finally rinsed with PBS and mounted on glass slides using Mowiol. Fluorescent images were taken with oil-immersion 63x magnification using a fluorescence microscope (Zeiss). Details of primary and secondary antibodies are given in Table S4.

### *In vivo* antitumor efficacy in subcutaneous MCF7 cell xenograft mouse model

Xenograft experiments were performed by WuXi AppTec (Hong Kong) Limited. The study was done in two phases. The first phase was a dose-response study in which two doses of each drug were selected based on the literature as being effective and tolerable (Table S1). In the second phase, low doses of each drug (around IC10 – IC20) were used at the beginning of the experiments and were further reduced throughout the study as indicated in Table S1. Experiments were performed in 6-8 weeks old female BALB/c nude mice weighing about 18-22 g. Three days before cell inoculation, 60-day release E2 pellets (Innovative Research of America) were implanted subcutaneously into the upper left flank. Then, 10 x 10^6^ MCF7 cells in 0.2 ml PBS + Matrigel (1:1) were inoculated subcutaneously into the right flank. One week later, when the average tumor volume reached 86 mm^3^, animals were randomized to groups of 5 using Microsoft Excel and treated with the corresponding drugs (Day 0). Tumor volume was measured twice every week using a caliper as follows: 0.5 x a x b^2^, in which a and b are the long and short diameters of the tumor, respectively. Drugs were dissolved in DMSO and then diluted by corn oil to the specified drug concentrations (final DMSO concentration was 10%). Drugs were administered daily by oral gavage either alone or in combination. Animals were checked every day for morbidity and mortality, and for signs of abnormal behavior such as changes in mobility, food and water consumption, BW, eye/hair matting, and any other effects due to the tumor or the drugs. BW was measured twice per week and in cases of BW loss >10%, animals were closely monitored, and if BW loss exceeded 15%, drug treatment was suspended, and animals were given estrogen-free nutrient gel to help their recovery. A limit of 20% BW loss was set as defined by the Institutional Animal Care and Use Committee (IACUC). The average tumor burden per mouse never exceeded 1,000 mm^3^. Manual bladder expression was performed for all mice. % change in tumor volume was calculated as follows: (Tumor volume on day “X” / Tumor volume on day “0” x 100) – 100, and % Change in BW was calculated as follows: (BW on day “X” / BW on day “0” x 100) – 100.

### Statistical analyses

Data were analyzed using MATLAB, GraphPad Prism 8.0.0 and Microsoft Excel. Unless otherwise indicated, differences between groups were analyzed with a one-way ANOVA test, error bars represent the standard deviation between replicates, and *p*-values ≤ 0.05 were considered statistically significant.

## Acknowledgments

We especially thank Pablo C. Echeverria for his grateful help and insights into the construction and analysis of PPI networks. We thank Prof. Leonardo Scapozza, and Santosh Anand for their help and fruitful discussions. We thank the team of WuXi AppTec for their highly collaborative and efficient services. We especially thank Chunxia Wang, Yajie Xiong, Zhengrong Yu, and Juergen Boer. We are grateful for all the gifts of reagents mentioned in the text.

## Funding

This work was supported by the Fondation Medic, the Fondation Boninchi, the Fondation Ernst et Lucie Schmidheiny, and the Canton de Genève.

## Author contributions

D.H. and D.P. conceived the study; D.H. designed and performed the *in silico* screen, most of the experiments, analyzed most of the data, prepared the figures and wrote the drafts; M.Z. contributed to the design and analysis of the TGMO-based screen; K.B. contributed to the FACS experiments; P.N.S. supervised the work and helped with the TGMO-based screen; D.P. supervised the work, curated the list of ERα interactors, contributed to the design of the experiments, wrote, and critically edited the manuscript. All authors have read and agreed to the published version of the manuscript.

## Competing interests

P.N.S. is an inventor of a patent on drug combination technology. Apart from that the authors declare that they have no competing interests.

## Data and materials availability

All data are available in the main text or the supplementary materials.

## The paper explained

### Problem

Most breast tumors are dependent on estrogen receptor α (ERα). Current therapy of breast cancer, for example with the anti-estrogen tamoxifen, is often limited by the development of drug-related resistance and toxicity. Therefore, there is an urgent need for new therapeutic approaches to overcome these problems. Multidrug combinations offer the advantage of lowering the doses and targeting the cancer from multiple angles, which can eventually lead to fewer adverse effects and reduced chances of drug resistance.

### Results

Using a computer-assisted approach to tap into multiple publicly available datasets, we established a list of drugs as candidates for synergistic drug combinations. Experimentally, we then identified two cocktails of three and four drugs that are efficient for a common subtype of ERα-positive breast cancer harboring PI3Kα mutations and have minimal effects on normal cells. These combinations also proved to be efficient for tamoxifen-resistant breast cancer cells and in relevant mouse models.

### Impact

The identified multidrug combinations with each drug given at very low doses have the potential to improve current breast cancer therapy. Owing to their enhanced safety profiles and increased efficacy for the treatment of both naive and drug-resistant ERα+ breast tumors, they may overcome many of the current clinical challenges.

## Notes

### Summary of Updates

Supplementary Material files have been included

