## Supplementary material for "Network-informed discovery of multidrug combinations for ERα+/HER2-/PI3Kα-mutant breast cancer": Supplementary Material.pdf

Hany et al.

**This file includes:**


Description of Tables S1 to S3
Tables S4 and S5
Description of the data files Data S1 to S5
Legends to Figures S1 to S10
Figures S1 to S10

**Description of Tables S1 to S3, which are available as separate files.**

**Table S1. Details and doses of the anticancer drugs used throughout the study.** The Microsoft Excel workbook contains information about the commercial source of the drugs, biological targets, drug development phase, physical properties, stock concentrations, and doses used for the TGMO-based screens, molecular assays, and xenograft experiments.

**Table S2. A list of predicted synergistic 2-, 3-, and 4- drug combinations from "Round 2" of the TGMO-based screen.** The Microsoft Excel workbook contains the list of candidate drug combinations for MCF7, MCF7-V, and MCF7/LCC2 cells, and their potential safety profile in MCF10A cells. Combinations of potential therapeutic relevance were selected for "Round 3" assays.

**Table S3. Status of *PIK3CA* gene mutations in different breast cancer cell lines.** The Microsoft Excel workbook contains the status of the tested mutations of the *PIK3CA* gene in exons 9 and 20 in the different breast cancer cell lines used in the study.

**Table S4. List of antibodies.**

| Antigen and/or antibody | Supplier | Catalog no. | Host | Clonality | Dilution |
|---|---|---|---|---|---|
| Caspase 3 | Cell Signaling Technology | 9662 | Rabbit | Polyclonal | 1:1,000 |
| Caspase 7, clone: (C7) | Cell Signaling Technology | 9494 | Mouse | Monoclonal | 1:1,000 |
| NFκB2 p100/p52 | Cell Signaling Technology | 4882 | Rabbit | Polyclonal | 1:1,000 |
| Phospho-Thr145 of p21 | Abbkine | ABP50378 | Rabbit | Polyclonal | 1:1,000 |
| p21 Cip1, clone: [GT1032] | GeneTex | GTX629543 | Mouse | Monoclonal | 1:1,000 for western blot and 1:500 for immuno-fluorescence |
| Rad51, clone [N1C2] | GeneTex | GTX100469 | Rabbit | Polyclonal | 1:1,000 |
| Lamin B1 | GeneTex | GTX103292 | Rabbit | Polyclonal | 1:1,000 |
| Phospho-Ser139 of histone γ-H2AX, clone: [GT1021] | GeneTex | GTX628996 | Mouse | Monoclonal | 1:2,000 for western blot and immuno-fluorescence |
| PARP (cleaved Asp214) | GeneTex | GTX132329 | Rabbit | Polyclonal | 1:1,000 |
| ERK 2, clone: (C-14) | Santa Cruz Biotechnology | Sc-154 | Rabbit | Polyclonal | 1:200 |
| Phospho-Tyr 204 of ERK, clone: (E-4) | Santa Cruz Biotechnology | Sc-7383 | Mouse | Monoclonal | 1:1,000 |
| Phospho-Tyrosine (P-Tyr-100) | Cell Signaling Technology | 9411 | Mouse | Monoclonal | 1:2,000 |
| a-tubulin, clone (DM1A) | Merck Millipore | CP06 | Mouse | Monoclonal | 1:5,000 |

| | | | | | |
|---|---|---|---|---|---|
| GAPDH, clone (6C5) | Abcam | ab8245 | Mouse | Monoclonal | 1:30,000 |
| Phospho-S311 of NFκB p65 | Santa Cruz | sc-135769 | Mouse | Monoclonal | 1:1,000 |
| Mouse IgG (H+L), HRP secondary antibody | Invitrogen | 31430 | Goat | Polyclonal | 1:10,000 |
| Rabbit IgG (H+L), HRP secondary antibody | Invitrogen | 31460 | Goat | Polyclonal | 1:10,000 |
| Mouse IgG (H+L), Alexa Fluor 488 | Invitrogen | A-11017 | Goat | Polyclonal | 1:5,000 |
| Mouse IgG (H+L), Alexa Fluor 546 | Invitrogen | A-11030 | Goat | Polyclonal | 1:1,000 |

**Table S5. List of specific primer sequences used for real-time RT-qPCR**

| **Gene** | **Forward 5'-3'** | **Reverse 5'-3'** |
|---|---|---|
| *VEGF* | CAGAAGGAGGAGGGCAGAATC | GTCCACCAGGGTCTCGATTG |
| *CDKN1A* | AGGTGGACCTGGAGACTCTCAG | TCCTCTTGGAGAAGATCAGCCG |
| *CDKN2A* | CTCGTGCTGATGCTACTGAGGA | GGTCGGCGCAGTTGGGCTCC |
| *CDKN1B* | ATAAGGAAGCGACCTGCAACCG | TTCTTGGGCGTCTGCTCCACAG |
| *TP53* | TAGTGTGGTGGTGCCCTATGA | ACACGCACCTCAAAGCTGTTC |
| *β-ACTIN* | CATGTACGTTGCTATCCAGGC | CTCCTTAATGTCACGCACGAT |

**Description of the Data files S1 to S5, which are available as separate files.**

**Data S1. ERα PPI network.** The Cytoscape file contains the network of primary PPIs of ERα with 322 protein interactors and secondary PPIs among them. Based on the analysis of network centrality measures, the top 50 interactors are highlighted in green.

**Data S2. List of ERα interactors.** The Microsoft Excel workbook contains the list of 322 ERα interactors that were given the feature "DP". The file was imported to Cytoscape to be integrated into the network of the human interactome.

**Data S3. Analysis of ERα PPI network centrality measures.** The Microsoft Excel workbook of the results of the network analysis done by Cytoscape on the ERα PPI. Results include coefficients for network centrality parameters, such as "Number of directed edges", "Radiality", "Stress", and others.

**Data S4. Genes co-expressed with ERα in ERα+ breast cancer datasets.** The Microsoft Excel workbook contains the co-expression coefficients obtained from datasets available from the online databases Oncomine (www.oncomine.org) and GOBO (http://co.bmc.lu.se/gobo).

**Data S5. Integrated network of SL between interactors of ERα, co-expressed, and frequently mutated genes in ERα+ breast cancer.** The Cytoscape file contains two networks; one represents SL between genes in breast cancer, and another network derived from the first one that contains only those genes that are ERα interactors, co-expressed, or frequently mutated genes in ERα+ breast cancer.

**Legends for Figures S1 to S10**

**Fig. S1. Frequently mutated genes and SL network in ERα+ breast cancer, and crosstalk between potential drug targets. (A,B)** Raw data of the most frequently mutated genes in ERα+ breast cancer obtained from the COSMIC (panel A) and IntOGen databases (panel B). In panel A, blue bars represent the number of samples with mutations and the red bars are the total number of samples analyzed. **(C)** Network of SL relationships in breast cancer. ERα interactors, genes co-expressed with ERα, and frequently mutated genes in ERα+ breast cancer are highlighted with color codes. Nodes represent genes and edges represent SL relationships between the connected genes. **(D)** Schematic illustration of relevant signaling pathways and the network of signaling crosstalk between them. Solid lines indicate a direct effect, whereas dotted lines indicate an indirect effect. The 9 drugs selected for "round 1" of the TGMO-based screen and their primary targets are included in the scheme. The scheme was created with Biorender.com based on information from the literature and the KEGG PATHWAY database.

**Fig. S2. Dose-response curves of the selected 9 drugs in cancer and non-cancer cell lines. (A-I)** Dose-response curves of the 9 selected drugs for "round 1" of the TGMO-based screen. The indicated cancer (blue and green) and non-cancer (orange) cell lines were treated with increasing doses of each drug and % ATP levels were calculated as the response. For each curve, a control group treated with vehicle was set to 100%. Data are represented as means ± SD (n = at least 3 independent samples).

**Fig. S3. Results of "Round 1" of the *in vitro* TGMO-based screen. (A-C)** Bar graphs of the % ATP levels of the breast cancer cell lines (blue) MCF7-V (panel A), MCF7 (panel B), and MCF7/LCC2 (panel C) treated with alpelisib (Alp), ulixertinib (Uli), obatoclax (Oba), talazoparib (Tala), MI-778 (MI-7), milciclib (Mil), UC2288 (UC2), C188-9 (C18), and OHT, tested alone or in combinations. Red bars show results obtained with the non-cancerous MCF10A cells. The data are presented as means ± SEM of 3 independent experiments, each with 3 independent replicates. The matrices below the graphs represent the scheme of the combinatorial design, in addition to the vehicle (DMSO) control in white, each drug was given at two dose levels, higher dose (dark gray) and lower dose (light gray). Values of the DMSO only controls were set to 100%.

**Fig. S4. Regression modeling of the results of "Round 1" of the *in vitro* TGMO-based screen. (A,B)** Estimated regression coefficients obtained by second-order linear regression of the calculated % ATP levels from "round 1" in MCF7-V (panel A) and MCF7/LCC2 (panel B) cells. Blue bars indicate regression coefficients of cancer cells, whereas red bars are show the TW as compared to MCF10A cells. Drugs highlighted in red in the x-axis labels represent drugs or combinations with unfavorable efficacy or toxicity profiles, which were therefore eliminated from "round 2". Statistical significance is shown as * for $p \leq 0.05$, ** for $p \leq 0.01$, *** for $p \leq 0.001$, and "ns" for non-significant results. Data are represented as means ± standard error of the means (SEM) of 3 independent experiments, each containing 3 independent replicates.

**Fig. S5. "Round 2" of the *in vitro* TGMO-based screen. (A-C)** Bar graphs of the % ATP levels of the breast cancer cell lines (blue) MCF7 (panel A), MCF7-V (panel B), and MCF7/LCC2 (panel C) treated with alpelisib (Alp), ulixertinib (Uli) (panels A and B), obatoclax (Oba), talazoparib (Tala) (panel C), MI-778 (MI-7), UC2288 (UC2), C188-9 (C18), and OHT, tested alone or in combinations. Red bars show results obtained with MCF10A cells. Note that the same values obtained with MCF10A cells were used in all three panels (A-C) for comparison. The data are presented as means ± SEM of 3 independent experiments, each with 3 independent replicates. The matrices below the graphs represent the scheme of the combinatorial design, in addition to the vehicle (DMSO) control in white, each drug was given at two dose levels, higher dose (dark gray) and lower dose (light gray). Values of the DMSO only controls were set to 100%. CIs calculated by CompuSyn are illustrated below the combinatorial matrices as a red color gradient. Low CI < 0.8 (light red) indicates synergistic combinations, whereas high CI > 1 (dark red) indicates antagonistic combinations. MT, monotherapy; Veh., DMSO control.

**Fig. S6. Regression modeling of the results of "Round 2" of the *in vitro* TGMO-based screen. (A-E)** Estimated regression coefficients obtained by second-order linear regression (panels A to C) or third-order linear regression (panels D and E) of the calculated % ATP levels from "round 2" in MCF7 (panel A), MCF7-V (panels B and D), and MCF7/LCC2 (panels C and E) cells. Blue bars indicate regression coefficients of cancer cells (panels A to E), whereas red bars show the TW as compared to MCF10A cells (panels A to C). For panel D, combinations highlighted in green in the x-axis labels have favorable efficacy profiles. Statistical significance is shown as * for $p \leq 0.05$, ** for $p \leq 0.01$, *** for $p \leq 0.001$, and "ns" for non-significant results. Data are represented as means ± SEM of 3 independent experiments, each with 3 independent replicates.

**Fig. S7. Specificity of the selected 7 drugs from "Round 2" in cancer cell lines representative of the different molecular subtypes of breast cancer. (A)** A scheme of the molecular subtypes of the cell lines used in "round 3" of the combinatorial screen based on ER$\alpha$ and HER2 expression. **(B-H)** Dose-response curves of the 7 selected drugs from "round 2" to be tested in different combinations in "round 3". The indicated breast cancer cell lines include ER$\alpha$+ (blue), ER$\alpha$+ but tamoxifen-tolerant or -resistant (green), and ER$\alpha$- (red) cell lines. Cells were treated with increasing doses of each drug and % ATP levels were calculated as the response. For each curve, a control group treated with vehicle was set to 100%. Data are represented as means ± SD (n = at least 3 independent samples). Note that the same data (curves) obtained with MCF7, MCF7-V, and MCF7/LCC2 cells were used as in Fig. S2, and plotted here for comparison.

**Fig. S8. "Round 3" of the *in vitro* TGMO-based screen. (A,B)** Heatmaps of the % ATP levels (blue gradient) of the breast cancer cell lines treated with alpelisib (Alp), ulixertinib (Uli), obatoclax (Oba), talazoparib (Tala), MI-778 (MI-7), UC2288 (UC2), and OHT, tested alone at two concentrations indicated as "lower dose" (panel A) and "higher dose" (panel B). Low % ATP levels (light blue) indicates high efficacy and high % ATP levels (dark blue) indicates low efficacy. The data are presented as means of 3 independent experiments. Values of DMSO control were set to 100%. **(C-**

**H)** Heatmaps of the % ATP levels (left side of the panel) of the predicted 2-drug (panels C and D), 3-drug (panels E and F), and 4-drug (panels G and H) combinations from "round 2". CIs calculated by CompuSyn are illustrated in the heatmap on the right of each panel (red gradient). Low CI < 0.8 (light red) indicates synergistic combinations, whereas high CI > 1 (dark red) indicates antagonistic combinations. Combinations were tested at lower (panels C, E, and G) and higher dose levels (panels D, F, and H). Red arrows indicate combinations with higher efficacy and synergy in ERα+ breast cancer cell lines and less toxicity in non-cancerous cells.

**Fig. S9. FACS analysis of PI/annexin V staining and evaluation of the effects of AUT on ERα, ERK, NFκB, and HIF1α activities. (A)** Representative dot plots of the flow cytometric analysis of MCF7-V cells double stained with PI/annexin V-FITC (AV-FITC). Cells were treated with alpelisib (Alp), UC2288 (UC2), or OHT alone, or in combinations. Populations of stained cells were gated into four quadrants: PI+/AV-FITC+ (upper right), PI-/AV-FITC+ (lower right), PI+/AV-FITC- (upper left), and PI-/AV-FITC- (lower left). **(B,C)** Luciferase reporter assays for the transcriptional activities of endogenous ERα in MCF7-V (panel B) and T47D (panel C) cells. Cells were transiently transfected with the ERE-Luc reporter plasmids. Relative luciferase activities (RLU) were normalized to the activities of the internal transfection standard Renilla luciferase. Cells were treated with alpelisib (Alp), UC2288 (UC2), and OHT, tested alone or in the 3-drug combination (red bars). Luciferase activities of the vehicle-treated control groups were set to 1. **(D)** Luciferase reporter assays for exogenously expressed ERα with HEK 293T cells. Cells were transiently co-transfected with the ERE-Luc reporter and either empty vector pSG5 (EV) or pHEG0 (ERα). Drug treatments were as indicated and mentioned in panels B and C. Relative luciferase activities (RLU) were normalized to the activities of the internal transfection standard Renilla luciferase. Luciferase activities of the control cells transfected with EV and treated with DMSO were set to 1. **(E)** Immunoblots of the indicated proteins tested in total cell lysates of MCF7-V cell line. Cells were treated for 24 hours with either alpelisib (Alp), UC2288 (UC2), OHT, or talazoparib (Tala) tested alone, or the 3-drug combination (Alp + UC2 + OHT), or the 4-drug combination (Alp + UC2 + OHT + Tala). EGF (10 ng/ml) was added to cells 15 min before harvesting to induce MAPK signaling. GAPDH was used as an internal standard. **(F,G)** Luciferase reporter assays for the transcriptional activities of endogenous HIF1$\alpha$ activity in T47D (panel F) and MCF7/TamR (panel G) cells. Cells were transiently transfected with the HRE-Luc reporter plasmids. Relative luciferase activities were calculated and cells were treated as in panel B and C. For bar graphs, data are represented as means ± SD (n = 4 independent samples for panels B and C, and n = at least 3 for panels D, F, and G). Statistical significance between the groups were analyzed by one-way ANOVA and *p*-values ≤ 0.05 were considered statistically significant. Drug doses are listed in Table S1.

**Fig. S10. Drug dose-responses with MCF7 xenografts. (A,B)** Line graphs of the % change in tumor volume (panel A) and % change in body weight (panel B) of MCF7 xenografts in Balb/c nude mice treated as indicated and as in Table S1. Data points represent the means and error bars are SEMs. (n = 5 mice / group at the start of the treatments). In the group treated with talazoparib 2 mg/kg, 1 mouse died and treatment was suspended for the rest of the group on day 9 due to severe toxicity (n = 0). Starting from day 13, 2 more mice died and 1 was euthanized due to poor conditions; only 1 mouse from this group remained alive till the end of the study but without being further treated as %BW loss was more than 15%. In the group treated with talazoparib 1 mg/kg, on day 6, treatment was suspended for one mouse (n = 4), on day 9, this mouse died and treatment was suspended for another one (n = 3), then on day 10, one more mouse died and treatment was suspended for the rest of the group (n = 0), and finally on day 12, the mice were euthanized due to severe poor conditions (n = 0). **(C)** Line graphs of the % change in body weight of MCF7 xenografts in Balb/c nude mice treated as indicated. Data points represent the means and error bars are SEMs. (n = 5 mice / group). For all panels, values on the first day of drug treatments (Day 0) were set to 0%. Differences in the mean values measured on day 28 of treatments were analyzed by unpaired student's t-test. *p*-values ≤ 0.05 were considered statistically significant. Drug doses are listed in Table S1.

Supplementary figure 1

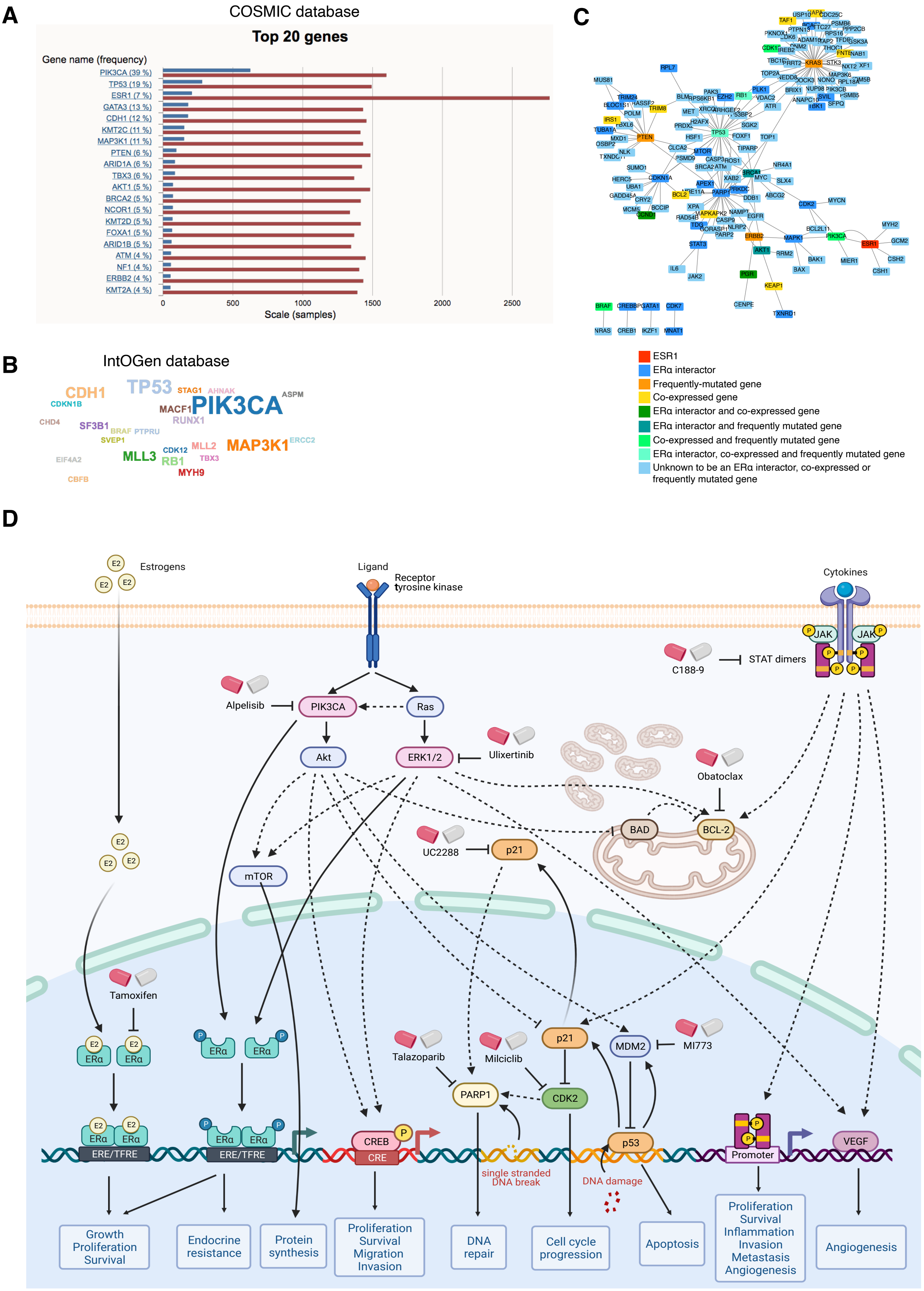

### Supplementary figure 2

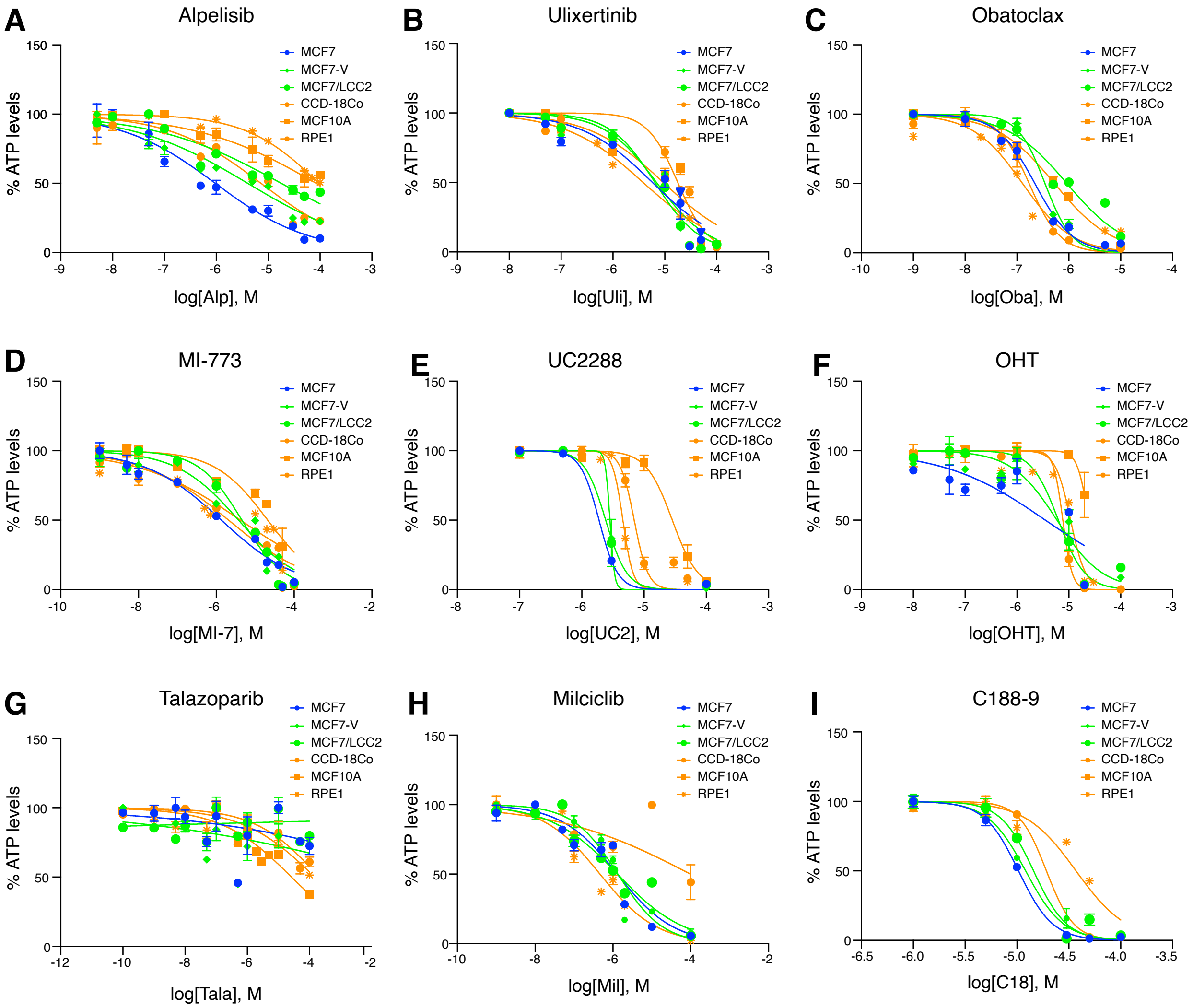

### Supplementary figure 3

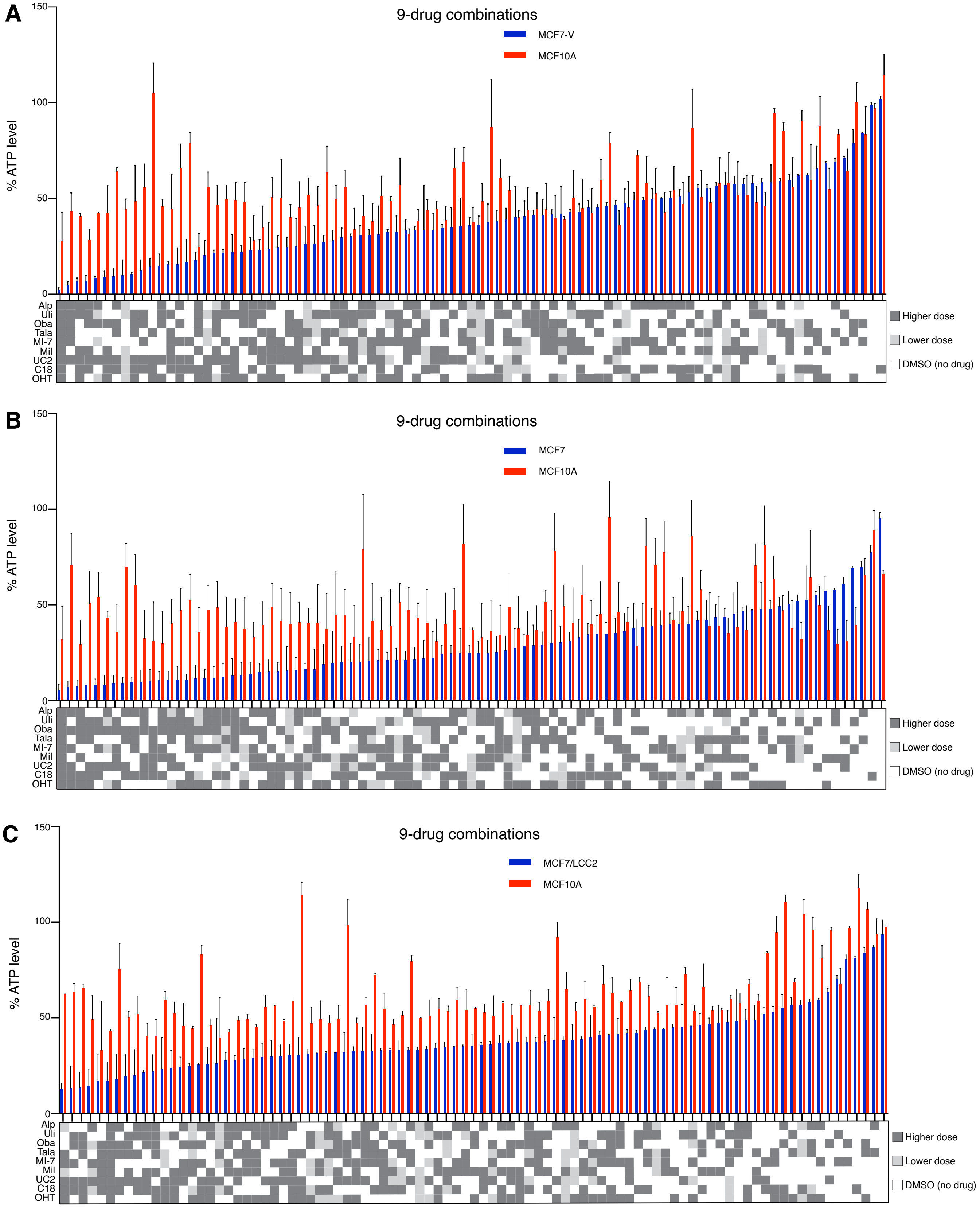

### Supplementary figure 4

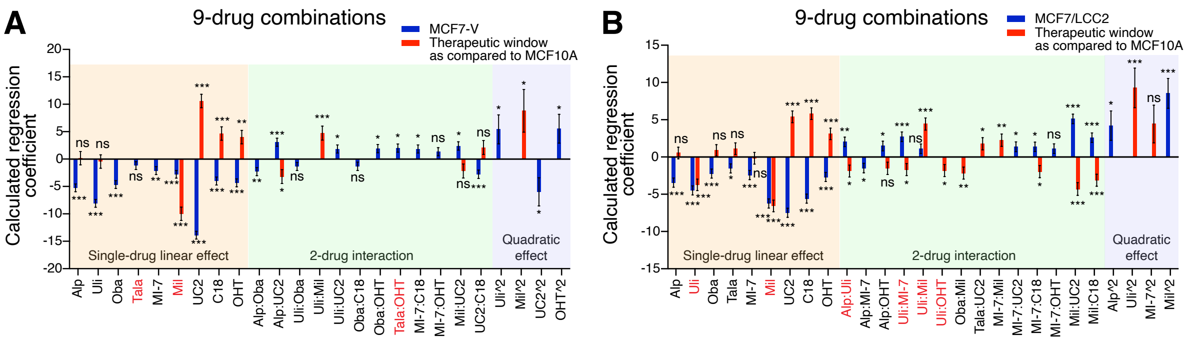

### Supplementary figure 5

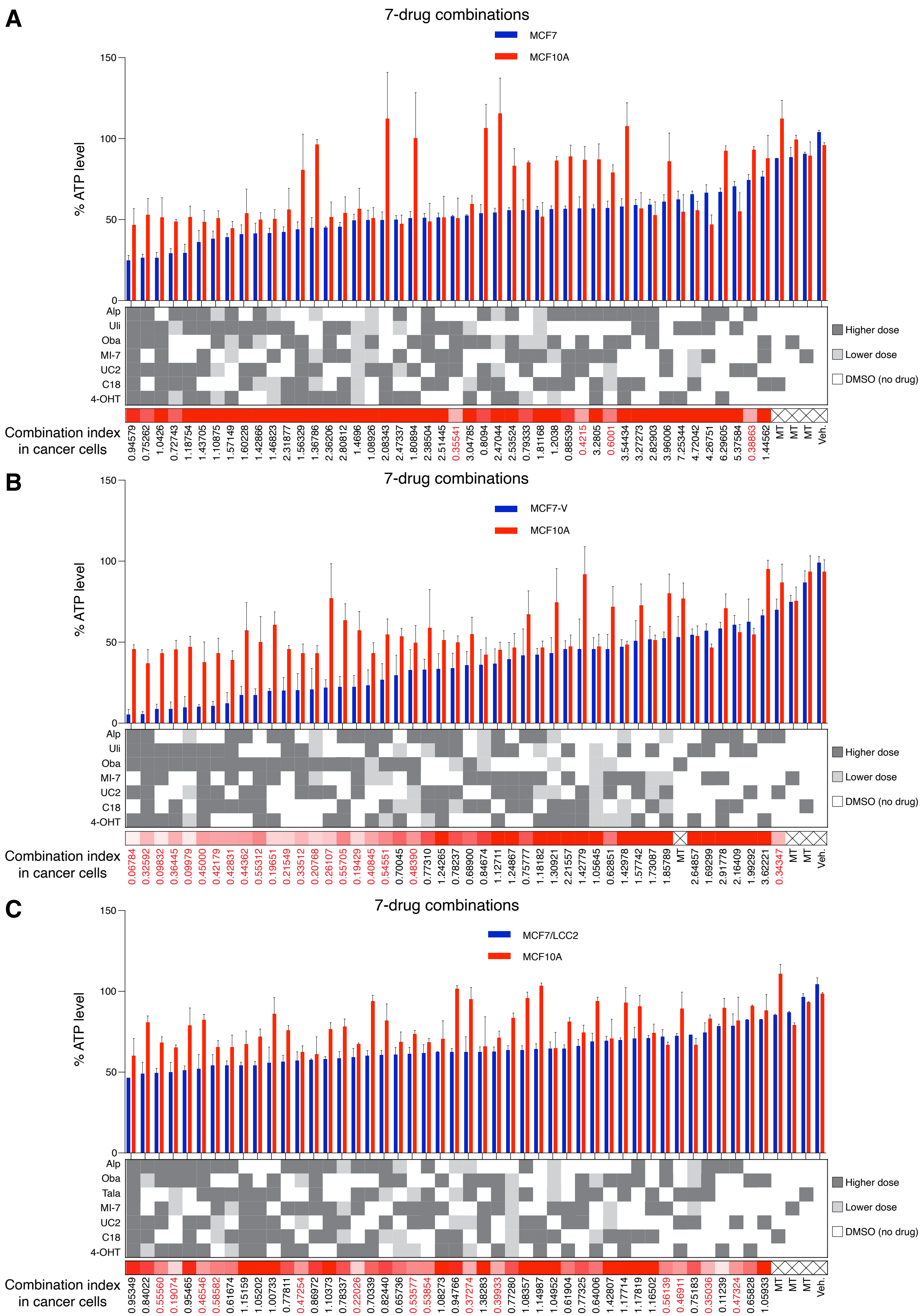

### Supplementary figure 6

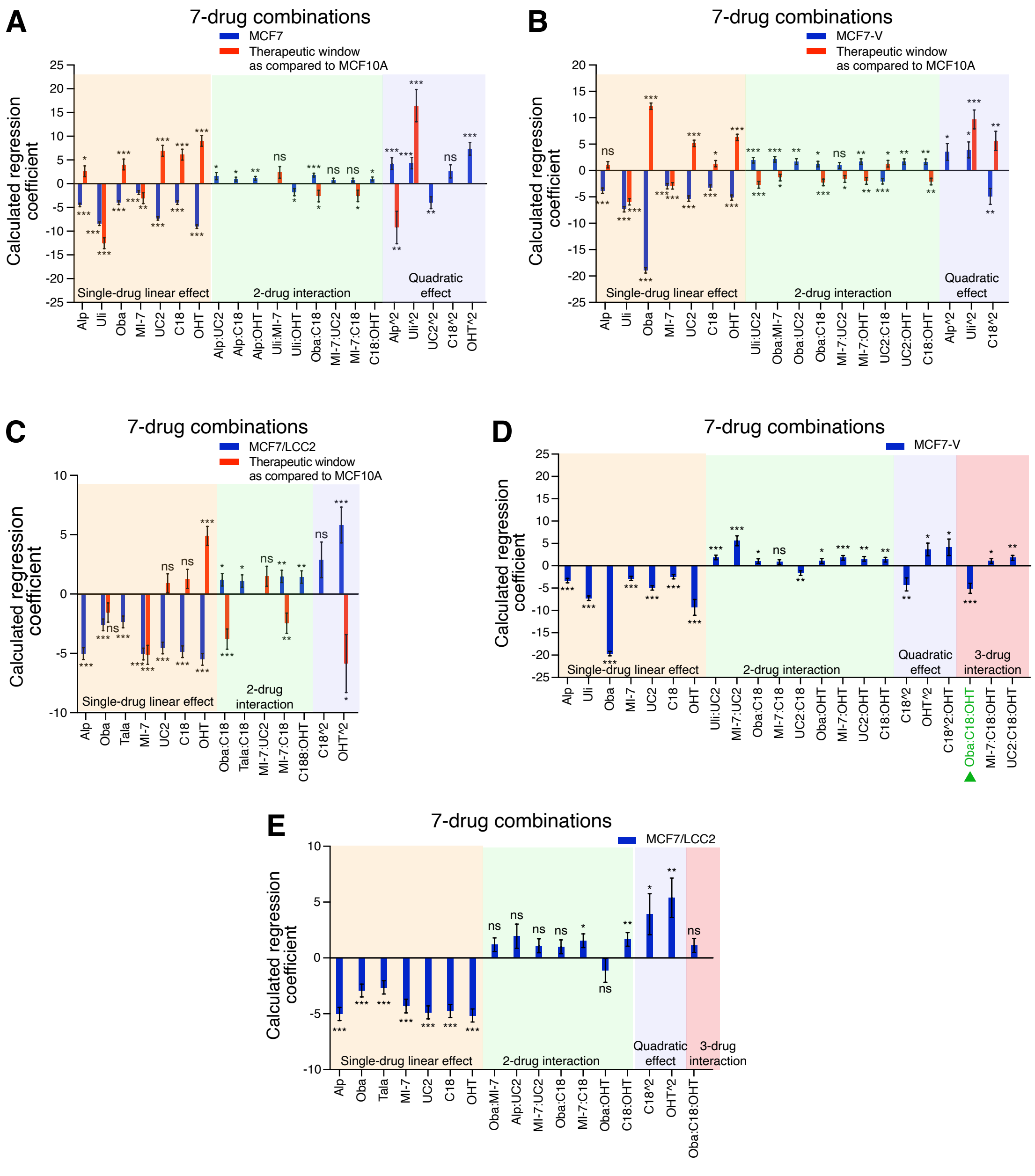

### Supplementary figure 7

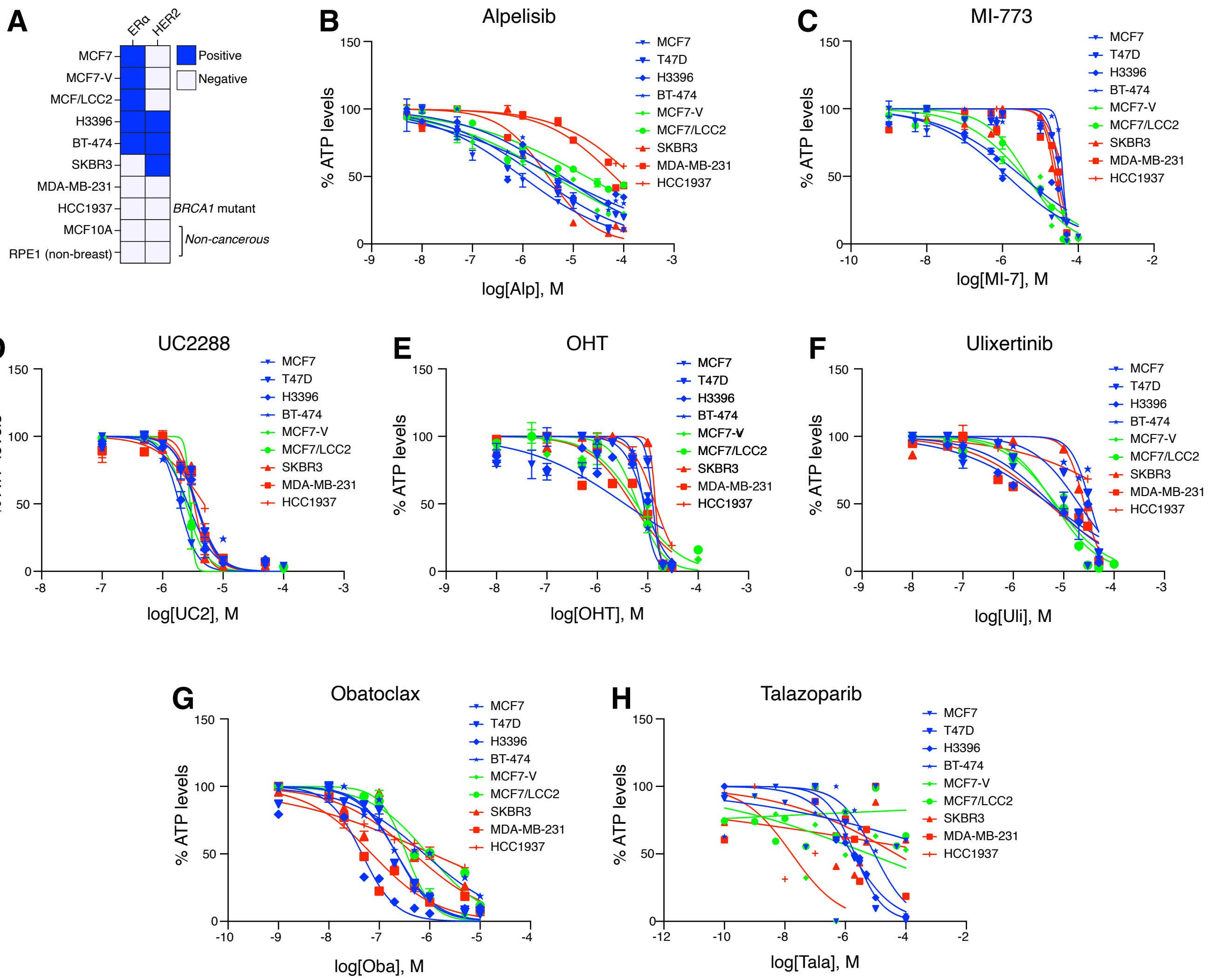

### Supplementary figure 8

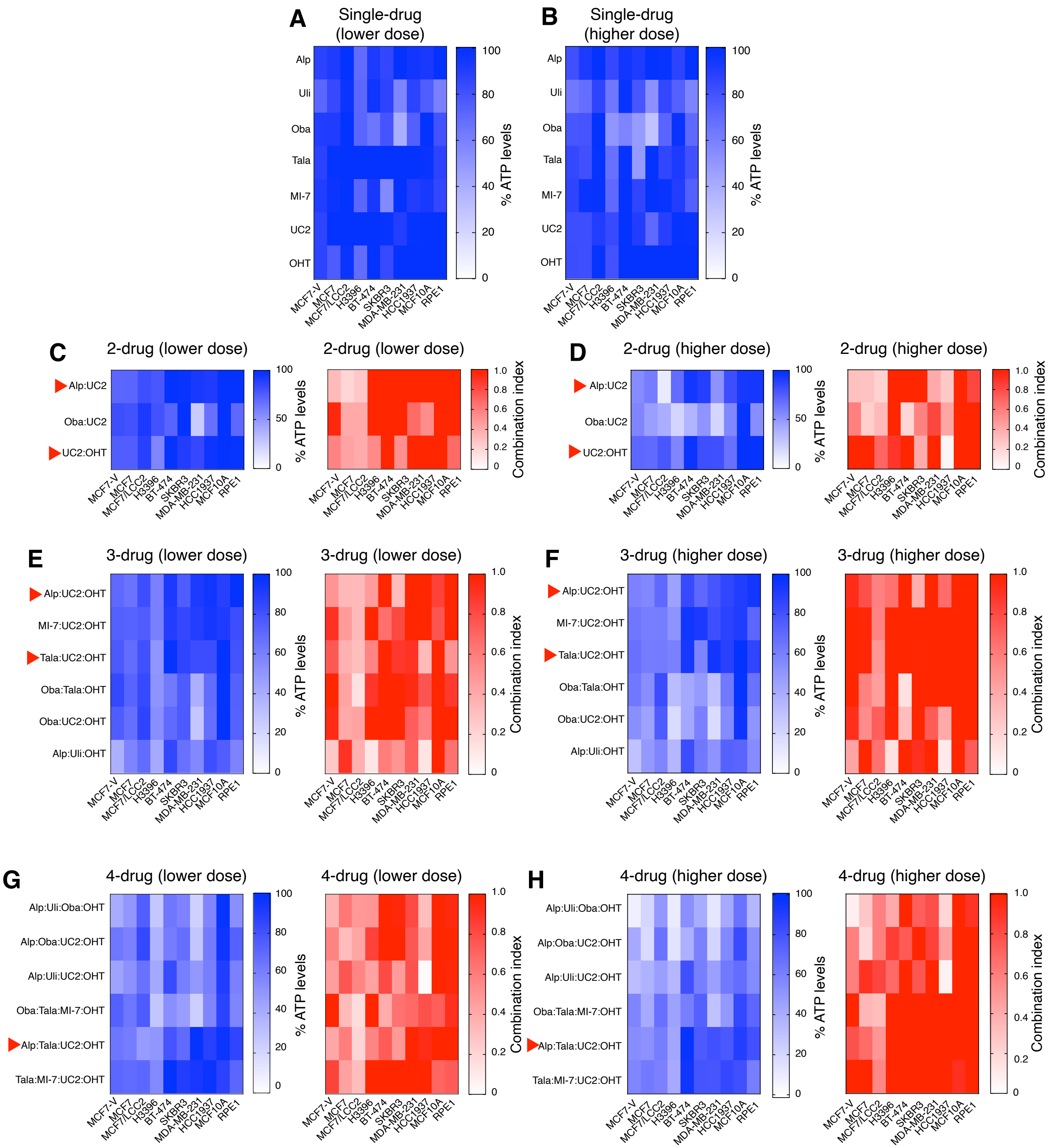

### Supplementary figure 9

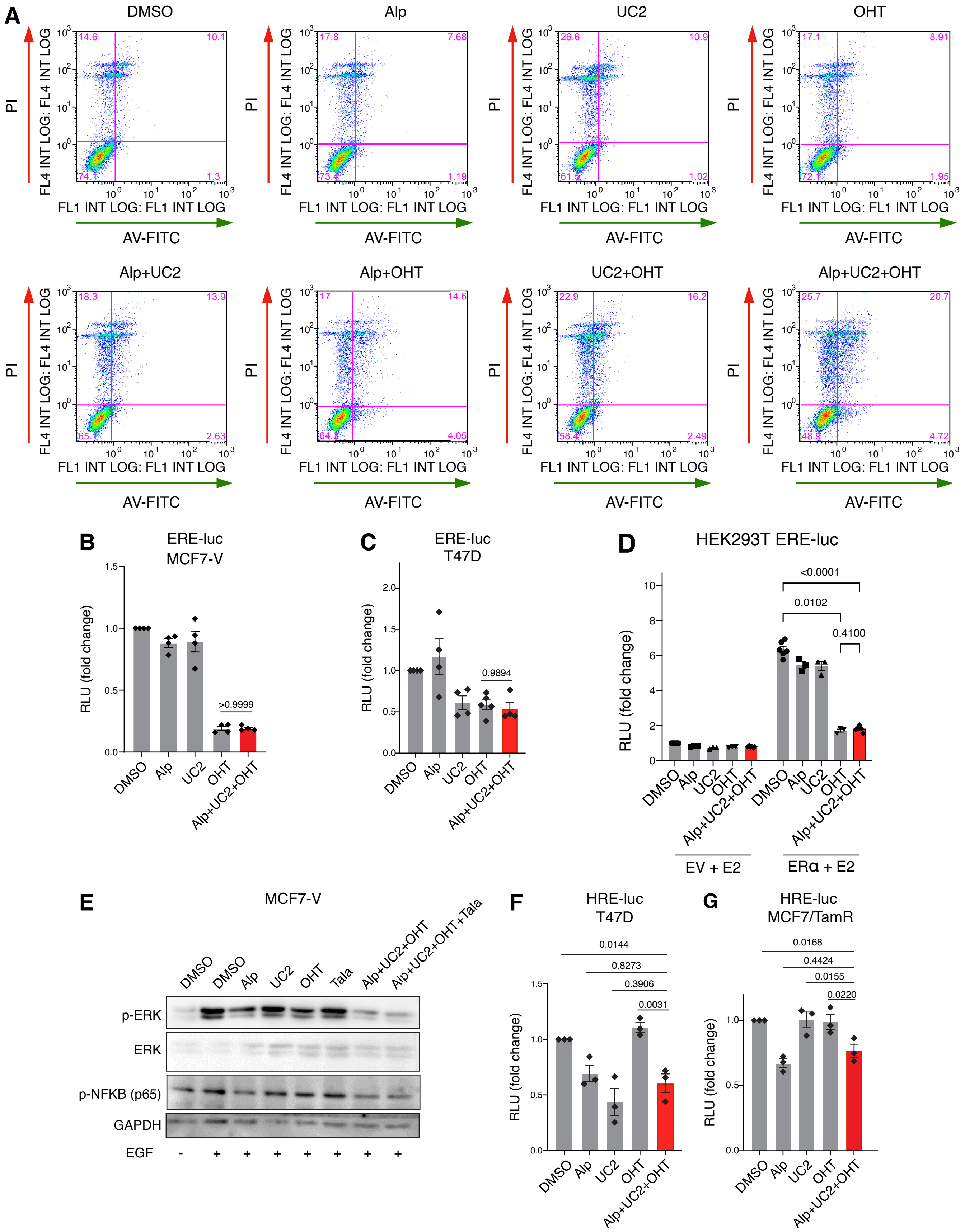

### Supplementary figure 10

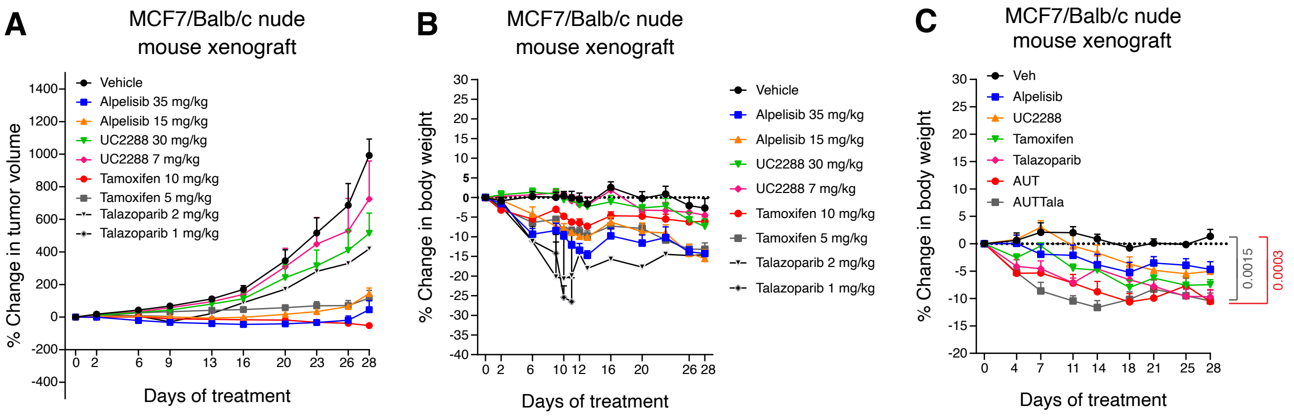
